# Nuclear envelope dysfunction drives premature aging and modulates heterochromatic methylome drift in *Arabidopsis*

**DOI:** 10.1101/2025.11.16.688749

**Authors:** Oscar Juez, Hidetoshi Saze

## Abstract

Aging involves progressive functional decline accompanied by molecular and epigenomic drift. Although nuclear-envelope (NE) defects cause premature aging in animals, the contribution of nuclear architecture to aging in plants remains unclear. Here we show that NE integrity is essential for maintaining epigenetic stability in *Arabidopsis thaliana*. Under short-day conditions, loss of *KAKU4* or *CRWN* proteins uncovers a photoperiod-sensitive, progeroid-like trajectory with premature senescence, transcriptional drift, and disruption of heterochromatin maintenance. Multi-omics analyses reveal that KAKU4 occupancy is confined to euchromatin and contracts with age, while CHH and CHG methylation erosion occurs within H3K9me2-enriched, transposon-dense heterochromatin. Wild-type plants normally exhibit age-associated CHH hypermethylation at transposable elements, a process abolished in NE mutants. Thus, nuclear-envelope integrity couples chromatin organization to the direction and rate of epigenomic drift, positioning perinuclear architecture as a conserved determinant of heterochromatin stability and aging trajectory across eukaryotes.

## Introduction

Aging is a conserved, multifactorial process that emerges across the tree of life and remains one of the central challenges in biology.^1,2^ Although the influential “hallmarks of aging” have been largely defined from animal studies (genomic instability, epigenetic alterations, stem-cell exhaustion, etc.), their universality beyond metazoans, particularly in modular organisms such as plants, remains unresolved.^3^ A growing body of evidence suggests that aging trajectories are constrained by conserved nuclear architectural features that predate multicellularity.^4,5^ Across eukaryotes, the nuclear envelope (NE) functions not merely as a physical boundary but as a dynamic interface for genome organization and transcriptional regulation.

Cytosine DNA methylation is a heritable, dynamic mark essential for genome stability and transcriptional control in animals and plants.^6^ In plants, methylation occurs in CG, CHG and CHH contexts (where H = A, T, or C), with CHH methylation at transposable elements (TEs) maintained by CHROMOMETHYLASEs (CMTs) and RNA-directed DNA methylation (RdDM) factors.^7,8^ Although plant methylomes are known to change with development and environmental stress, the directionality, chromatin-state specificity, and mechanistic basis of age-associated methylation drift in plants remain poorly understand.

Nuclear architecture is a strong candidate regulator of methylome stability. In animals, defects in nuclear lamina components cause progeroid syndromes and lead to accelerated methylome erosion, particularly within lamina-associated, late-replicating heterochromatin.^9–11^ While plants lack canonical lamins, thy form a lamina-like scaffold composed of Nuclear Matrix Constituent Protein (NMCP)/ CROWDED NUCLEI (CRWN) proteins and inner-nuclear-membrane factors such as KAKU4 and MAN1, which collectively tether and organize chromatin at the nuclear periphery.^12–16^ Whether plant NE integrity in plants constrains chromatin-state stability and shapes aging-associated methylome patterns in vivo has remained an open question.

Here, we address this gap by integrating lifespan phenotyping, transcriptomics, ChIP-seq and Enzymatic Methyl-seq (EM-seq) in *Arabidopsis thaliana* NE mutants. First, we demonstrate that loss of KAKU4 (and selected CRWN paralogs) triggers a photoperiod-sensitive, progeroid-like trajectory under short day conditions—characterized by premature leaf senescence, meristem failure, and shortened vegetative lifespan—without simply accelerating developmental phase transitions. Transcriptome profiling reveals a shift along the wild-type aging axis, marked by activation of stress/senescence pathways while down-regulating RNA-biosynthetic genes, linking nuclear architecture to age-aligned transcriptional drift. Second, chromatin-state-resolved methylome analysis show that in wild-type, CHH methylation progressively accumulates with age within TE-rich pericentromeric heterochromatin, consistent with reinforced TE silencing. In contrast, severe NE mutants fail to maintain this age-linked CHH methylation reinforcement, exhibiting targeted erosion of CHH and CHG methylation across heterochromatin. Notably, KAKU4 occupancy is biased toward euchromatin—enriched at open chromatin regions such as gene promoters and depleted over TE bodies—indicating that heterochromatic methylation failure arises indirectly through disrupted perinuclear silencing environments rather than through the loss of direct heterochromatin tethering. Together, these findings position NE integrity as an upstream determinant of heterochromatin maintenance and reveal that CHH methylation stability is architecture-dependent. This establishes a mechanistic bridge between nuclear-envelope integrity, chromatin-state stability, and age-associated methylome drift in plants, situating our work at the intersection of nuclear architecture and epigenetic aging.

## Results

### A progeroid-like premature aging in *Arabidopsis* nuclear-envelope mutants

In *Arabidopsis thaliana*, aging is closely intertwined with developmental progression because photoperiodic induction of flowering triggers whole-plant senescence on a compressed timescale.^17,18^ To disentangle chronological aging from developmental phase, we exploited photoperiod sensitivity of *Arabidopsis* by growing nuclear-envelope (NE) mutants (*CRWN1-4*, *KAKU4*) and photoperiod-response control mutants under short-day (SD; 8 h light/16 h dark) conditions, which extends the vegetative lifespan, and long-day (LD; 16 h light/8 h dark) conditions, which accelerate flowering. We asked whether disruption of NE integrity reveals an intrinsic, time-dependent senescence that is otherwise obscured when lifespan is developmentally compressed.

Under LD, all genotypes, including *kaku4* and *crwn* mutants, bolted and senesced within ∼55 days after sowing (DAS) without clear differences in lifespan or developmental trajectory (Supplementary. Figs. 1 and 2A) in our growth condition, consistent across replicates but distinct from a previous report.^19^ In contrast, SD growth condition exposed a progressive, degenerative sequence of senescence in *kaku4* and *crwn* genotypes (including *crwn1/2*, *crwn1/4*, *crwn2/3* double mutants) that was absent in wild-type plants (Fig. 1A, B; Supplementary Fig. 2B, C). Senescence initiated as basal-to-apical chlorosis and necrosis of the oldest rosette leaves, followed by a gradual loss of photosynthetic area and eventual rosette collapse. The process culminated in loss of meristematic competence, frequently manifesting as bolting failure or post-initiation arrest, despite continued chronological aging (Fig. 1B). Time-course imaging of *kaku4* rosettes at 90, 115, and 160 DAS under SD captured the sequential tissue degeneration and aborted meristems, accompanied by the retention of small juvenile leaves, indicating a defect in maintenance rather than an altered developmental transition (Fig. 1B).

**Figure 1.**
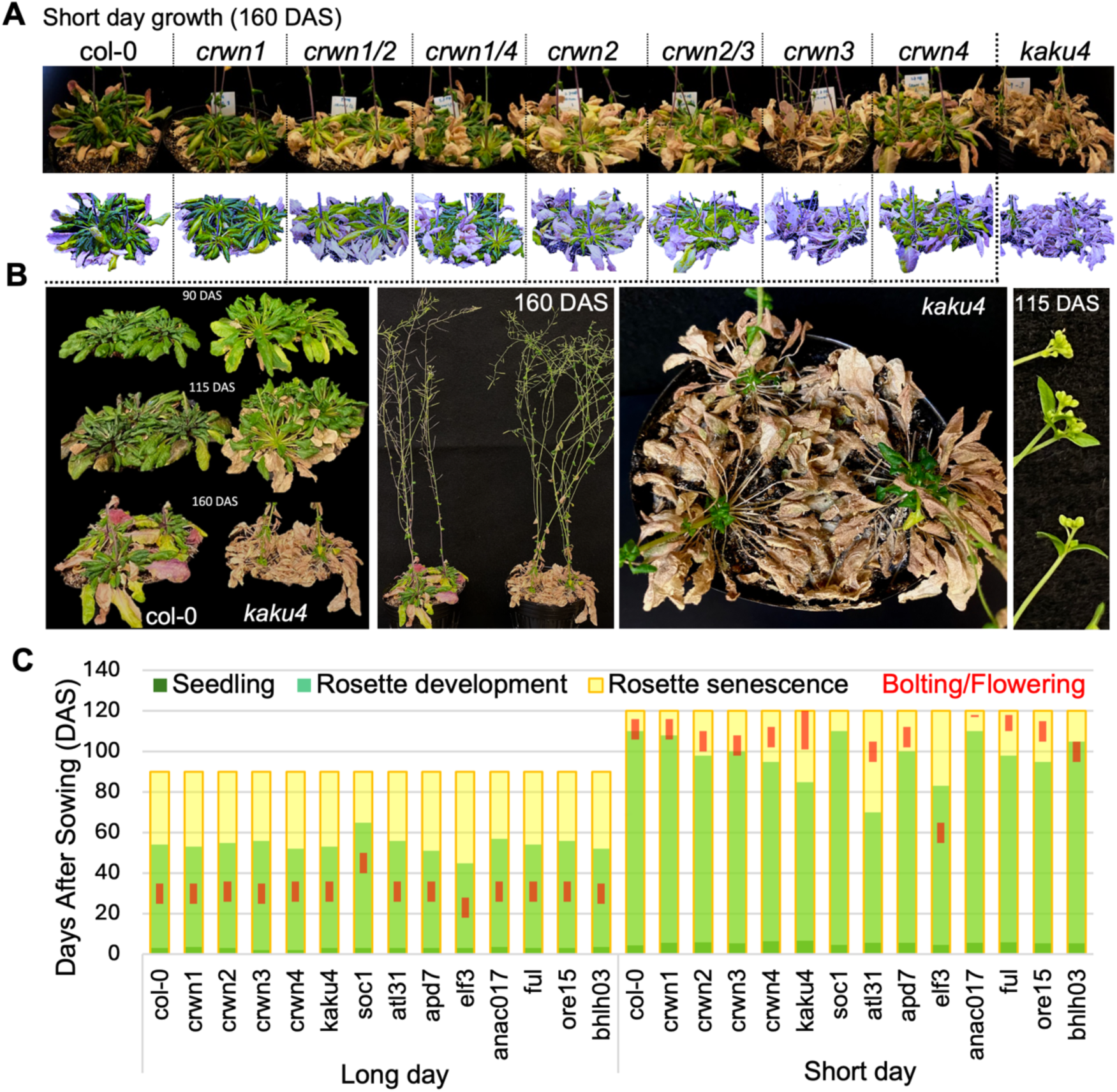
CRWNs and KAKU4 mutants exhibit photoperiod-dependent premature aging and shortened vegetative lifespan. **(A)** Whole-plant and rosette phenotypes of Arabidopsis *kaku4* and *crwn1-4* single and double mutants grown under short-day (SD; 8 h light) conditions for 160 DAS. Under SD, mutants exhibit a progressive progeroid phenotype characterized by basal leaf necrosis, loss of meristematic competence, and bolting arrest. Bottom: color-adjusted rosette projections to enhance visualization of senescent tissue (highlighted in purple). **(B)** Time-lapse images of kaku4 rosettes at 90, 115, and 160 DAS under SD, highlighting progressive tissue collapse, floral arrest, and emergence of juvenile traits (persistent small leaves). Right panels show flowering defect, arrested or incompletely elongated inflorescences. **(C)** Quantitative lifespan analysis of T-DNA insertion lines in nuclear envelope genes under LD (16 h light, 90 DAS) and SD (8 h light, 160 DAS). Lifespan was partitioned into seedling (dark green), rosette development (light green), rosette senescence (yellow), and bolting/flowering (red) phases. Each bar represents the mean of 9 plants (3 per pot × 3 pots). Under SD, *crwn3, crwn4*, and *kaku4* mutants specially exhibited precocious senescence and reduced vegetative lifespan. Internal controls (e.g., *atl31*, *elf3*, *soc1*, *ful*) confirm photoperiod-sensitive modulation of lifespan dynamics.

**Figure 2.**
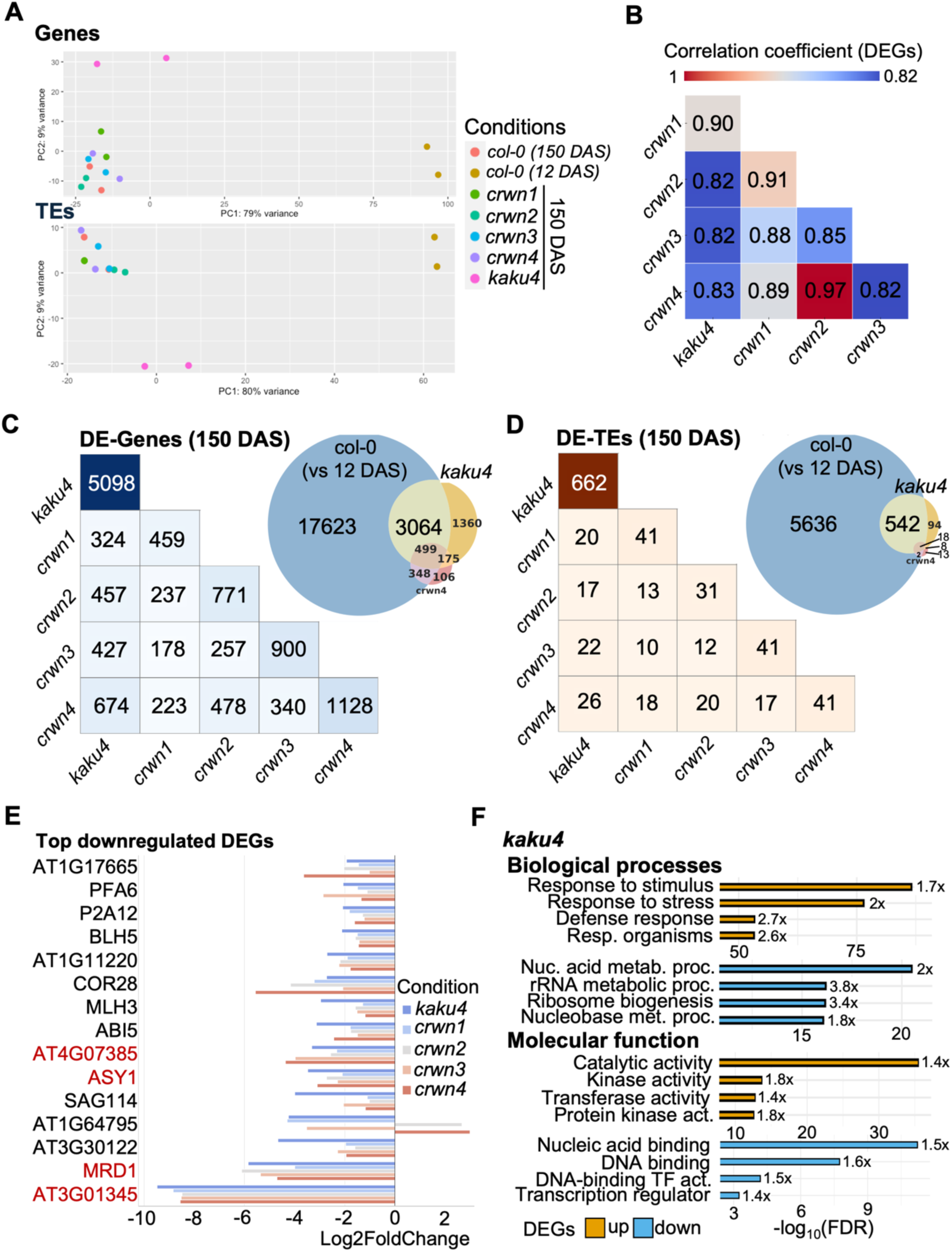
Transcriptomic drift in nuclear envelope mutants disrupts genes, TEs, RNA biosynthesis pathways. **(A)** Principal component analysis (PCA) of RNA-seq expression profiles in *Arabidopsis* wild-type (Col-0, 12 DAS and 150 DAS) and nuclear-envelope mutants (*kaku4*, *crwn1–4* at 150 DAS). Top: PCA of protein-coding genes; bottom: PCA of transposable element (TE) expression. Expression divergence along PC1 (genes: 79%, TEs: 80%) separates young vs aged wild-type and distinguishes *kaku4* from *crwns*. **(B)** Spearman correlation matrix of DESeq2-normalized gene expression profiles across samples. Values represent pairwise correlation coefficients among genotypes. *kaku4* shows the strongest divergence from aged wild-type, with lower correlation to both *crwn* mutants and Col-0. **(C)** Overlap of differentially expressed genes (DEGs) and (D) transposable elements (DETEs) across nuclear-envelope mutants under SD. Left: pairwise overlap matrices showing the number of shared DEGs/DETEs compared to control Col-0 at 150 DAS. Right: scaled Venn diagrams comparing DEGs/DETEs in *kaku4* and *crwn4* to aging-associated expression in Col-0 (150 DAS vs 12 DAS). Venn areas are proportional to overlap. All features are significant (DESeq2, padj < 0.05). **(E)** Common downregulated DEGs across nuclear-envelope mutants against Col-0 at 150 DAS. Horizontal bar plots show log₂ fold-change values of the top 25 up- and down-regulated transcripts shared between *kaku4* and *crwn1-4*. In red, lowest DEGs correspond to lncRNAs or 24-nt siRNAs islands, chromatin regulators, or senescence-linked loci (e.g., MRD1, ASY1). Expanded in Sup. Fig. 6A. **(F)** Functional enrichment of DEGs in *kaku4*. GO terms enriched among upregulated (gold) and downregulated (blue) genes were identified using the PANTHER Overrepresentation Test with FDR correction. Bar plots show -log₁₀(FDR) values and corresponding fold-enrichment for selected non-redundant categories from biological process, and molecular function. Expanded in Sup. Fig. 7-9.

To quantify these trajectories, we partitioned the lifespan into seedling, rosette development, rosette senescence, and bolting phases, and scores nine plants per genotype (three per pot, three pots) under both LD and SD conditions (Fig. 1C; Supplementary Fig. 1). Under SD, *kaku4*, *crwn1-4* mutants initiated rosette senescence significantly earlier (shorter time to first yellowing), spent a greater proportion of lifespan in the senescence phase, and exhibited markedly reduced bolting frequency by 120 DAS. When bolting occurred, inflorescences were frequently arrested or incompletely elongated. These differences were not apparent under LD, confirming that photoperiodic compression of vegetative lifespan conceals an underlying progeroid-like trajectory.

To assess the observed SD-specific senescence in NE mutants, we examined a panel of control mutants affecting photoperiod integration, metabolic signaling, and lifespan regulation (Fig. 1C, Supplementary Fig. 3): Photoperiod integrators: *atl31* (carbon/nitrogen-responsive E3 ligase) displayed accelerated senescence under SD but retained meristematic competence and bolting capacity,^20^ whereas *elf3* (an evening complex repressor of *GI* and *CO*) bolted early under both LD and SD with extreme stem elongation;^21^ neither recapitulated the *kaku4/crwn*-like senescence. Metabolic and hormonal pathways: *anac017* (mitochondrial retrograde NAC) and *apd7* (auxin-related clade-D PP2C) moderately shortened lifespan under LD by accelerating reproductive transition (*anac017*)^22^ or inducing vegetative collapse (*apd7*),^23^ but neither showed arrested meristem. Developmental-phase regulation: the *soc1* mutant delayed senescence under LD (as defined in Methods) yet remained fully vegetative state under SD even at 115 DAS, confirming that extended vegetative duration alone does not trigger senescence-driven collapse.^24,25^ Within the NE network, *man1* mutants also exhibited pronounced premature aging similar to *kaku4*, whereas *Plant Nuclear Envelope Transmembrane protein 2* (*pnet2*) single mutant maintained vegetative integrity and reproductive competence under SD (Supplementary Fig. 3D).^26^ These comparisons indicate that premature senescence is specific to particular NE components rather than a general consequence of NE perturbation. Together, these results demonstrate that prolonged vegetative lifespan under SD condition unmasks a latent, photoperiod-sensitive senescence program in *kaku4* and *crwn* mutants, revealing a progeroid-like premature aging phenotype in plants caused by NE dysfunction.

### Global transcriptomic drift in nuclear-envelope mutants disrupts transcriptional transitions during aging and misregulates defense and RNA-biosynthesis pathways

To assess how NE dysfunction affects transcriptional transition during aging, we compared transcriptomes of NE mutants with those of wild-type (Col-0) at defined chronological stages (12 and 150 DAS) under the SD condition. Principal Component analysis (PCA) revealed extensive variance between aged (150 DAS) and young (12 DAS) wild-type plants, explaining 79% (PC1) and 9% (PC2) of total variance for genes, and 80% and 9% for transposable elements (TEs) (Fig. 2A). Among the mutants, *kaku4* exhibited the largest displacement along PC1, positioned beyond aged Col-0 for both genes and TEs, whereas *crwn* single mutants clustered closer to aged wild-type. Expression distributions were consistent among biological replicates, particularly in young seedlings, with greater heterogeneity in older sample-most notably in *kaku4* (Supplementary Fig. 4A, B; see Methods). Pairwise Spearman correlations and Euclidean distance of global differential gene and TE expression profiles relative to aged Col-0 further supported these trends (Fig. 2B, Supplementary Fig. 4C). Inter-CRWN correlations were uniformly high (e.g., *crwn2-crwn4*, ρ ≈ 0.97), whereas *kaku4* showed the lowest similarity to other NE mutants (ρ ≈ 0.82-0.83), suggesting the activation of distinct, mutant-specific gene sets that are not typically engaged during developmental aging.

Differential expression analysis (DESeq2, *padj* < 0.05) revealed that at 150 DAS, compared to Col-0, *kaku4* displayed extensive transcriptional remodeling with ∼5,098 differentially expressed genes (DEGs), far exceeding that of CRWN genotype (459-1,128 DEGs; Fig. 2C, left). Likewise, *kaku4* exhibited the highest number of differentially expressed TEs (DETEs; ∼662) compared with *crwn* mutants (31-41 DETEs; Fig. 2D, left). When compared with the wild-type aging transcriptome (150 DAS vs. 12 DAS), *kaku4* shared a large subset of DEGs (∼3,064) and DETEs (∼542), but also possessed a substantial number of uniquely misregulated genes (∼1,360 DEGs and ∼94 DETEs) (Fig. 2C-D, right; Supplementary Fig. 4A,B). These results indicate that *kaku4* exaggerates normal age-associated transcriptional drift while also inducing unique, architecture-specific misregulation.

We further extracted directionally consistent DEGs shared between *kaku4* and *crwn1-4* mutants under SD (relative to Col-0, 150 DAS; Fig. 2E, Supplementary Fig. 5, 6). The shared gene set included several methylation-sensitive loci such as *AT3G01345*,^27^ as well as non-coding RNA genes *Mto1 Responding Down 1* (MRD1) and *RNA In The Antisense orientation* (*RITA*)/*AT1G64795*.^28^ Functional enrichment analysis of DEGs across mutants (Fig. 2F, Supplementary Fig. 7-9) revealed two major trends: (1) Upregulated genes were strongly enriched for biological processes related to defense and stress-response pathways, and (2) Downregulated genes were enriched for RNA biosynthesis pathways. The coordinated activation of defense-related genes aligns with previous reports in *crwn* mutants,^29,30^ reinforcing a conserved role of NE proteins in modulating stress and immune responses in *Arabidopsis*. Collectively, these findings indicate that disruption of nuclear-envelope architecture induces a global transcriptomic drift that both accelerates age-aligned expression transitions and distorts regulatory programs controlling defense and RNA biosynthetic functions.

### KAKU4 occupancy landscape reveals euchromatic preference and age-dependent contraction

To map KAKU4-chromatin interactions, we performed ChIP-seq using *Arabidopsis* line expressing KAKU4-eYFP (Fig. 3A, Supplementary Fig. 10A)^15^ at 10 DAS seedling and 120 DAS senescent rosette stages under SD conditions. In seedlings, a broad-peak calling (q < 0.05, cutoff 0.1) identified 4,461, 3,205, and 3,963 peaks across three biological replicates (mean = 3,876 ± 645 SD; Fig. 3B, Supplementary Fig. 10B). In senescent rosettes, however, only 381, 861, and 293 peaks per replicate were detected (mean = 512 ± 296 SD), representing an approximately 7.6-fold reduction relative to seedlings (*p* = 0.00115, two-sided t-test; Fig. 3B). Annotation of peak locations revealed that KAKU4 binding in seedlings was overwhelmingly gene-proximal: ∼60% of peaks overlapped promoters and transcription start site (TSS) regions, and ∼15% occurred within exons, with only minor fractions mapping to introns or distal intergenic regions (∼4-7%; Fig. 3C). By 120 DAS, this composition shifted markedly—promoter-associated peaks dropped to ∼31%, and TSS-associated peaks to ∼17% of the total—indicating a redistribution and contraction of KAKU4 binding away from promoter and TSS regions during aging.

**Figure 3.**
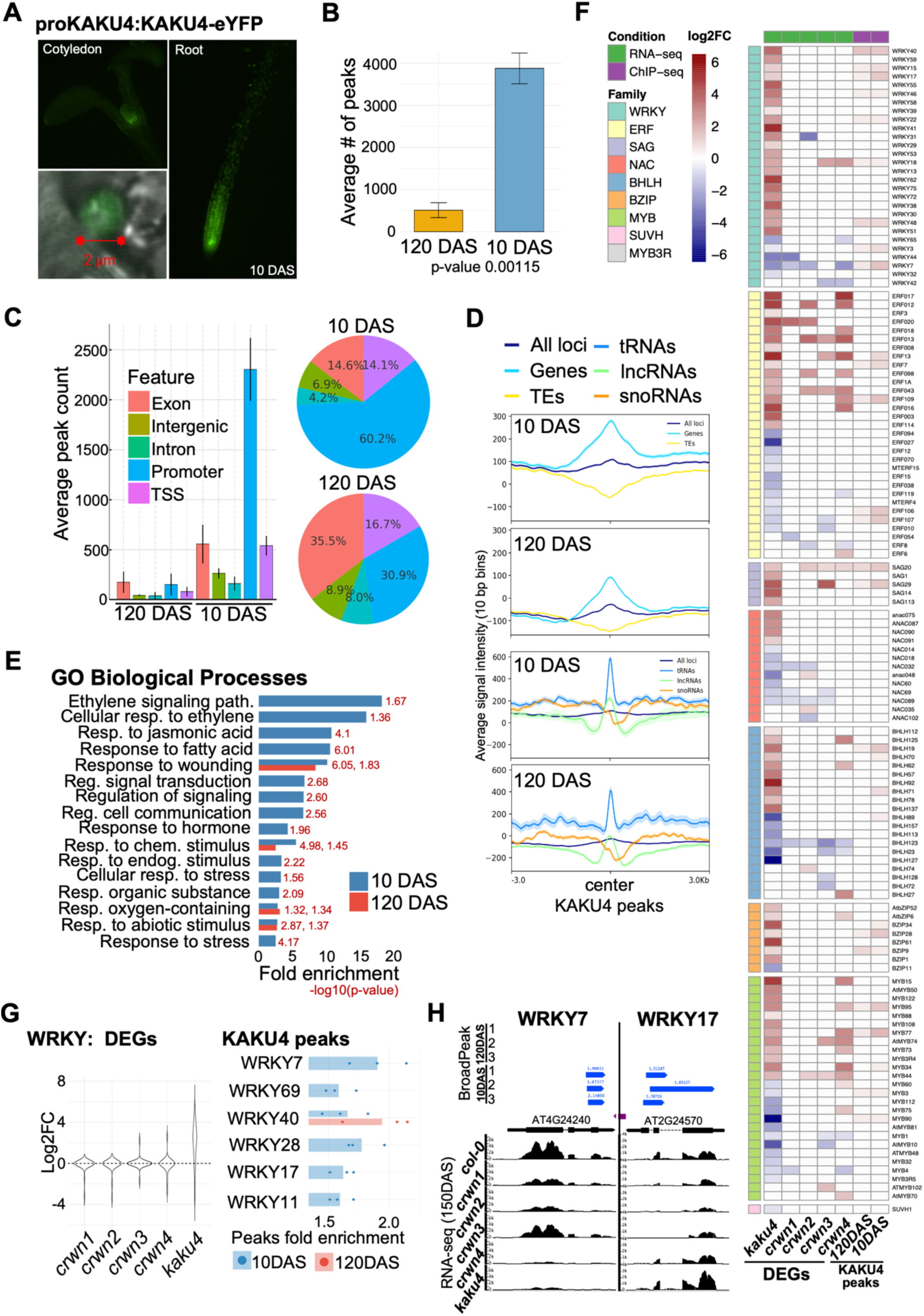
KAKU4 preferentially associates to euchromatic features that redistributed during aging. **(A)** Subnuclear localization of KAKU4–eYFP fusion protein in 10-day-old seedlings. Confocal images of proKAKU4:KAKU4-eYFP show nuclear localization in cotyledon and root epidermal cells. Bottom: magnified nucleus highlights peripheral ring signal. Scale bar = 2 μm. Expanded in Sup. Fig. 10A. **(B)** Total number of MACS2-called broad peaks (q < 0.05, –broad-cutoff 0.1) in KAKU4–EYFP ChIP-seq from 10 DAS and 120 DAS tissues. Two-sided unpaired Welch’s t-tests; bar charts show mean ± SD (n = 3). Expanded in Sup. Fig. 10B. **(C)** Distribution of KAKU4-bound peaks across genomic features. Left: barplot shows number of peaks overlapping exons, introns, promoters, TSSs, and intergenic regions. Right: pie charts display relative proportions for 10 DAS and 120 DAS samples, based on HOMER annotations. **(D)** Metaprofiles of all loci, genes, TEs, tRNAs, lncRNAs, snoRNAs intercepted to KAKU4-eYFP peaks. Signal centered on high-confidence KAKU4 peaks (10 bp window, ±3 kb from summit). **(E)** Gene ontology enrichment for genes with age-biased KAKU4 peaks. Barplots show – log10(FDR) *p*-values for GO biological process terms enriched among young- or old-biased peaks (Fisher’s exact test, BH correction). Blue bars = young-enriched; red = old-enriched. **(F)** Heatmap of RNA-seq and ChIP-seq log₂ fold-changes for age-related TF families. Each row corresponds to a WRKY, NAC, ERF, MYB, BZIP, or SAG gene with differential expression (left DEGs columns; at 150 DAS vs Col-0) across nuclear-envelope mutant (*kaku4*, *crwn1-4*). Last two columns (right); KAKU4-eYFP ChIP-seq normalized mean log₂FC (n = 3) signals for 120 and 10 DAS plants. **(G)** Left; differential gene expression across WRKY genes in panel F, shown as the combined distribution of expression per condition. Right; bars show the mean MACS2 fold-enrichment (IP over input; linear scale) for peaks assigned to each gene, averaged across three biological replicates within each age group. Points indicate per-replicate means. **(H)** Genome browser views of representative KAKU4-bound genes *AT4G24240* and *AT2G24570*. ChIP-seq tracks (3 replicates per age), MACS2 peak calls, and RNA-seq coverage of mutants are shown. Reduced KAKU4 occupancy in aged samples coincides with expression changes in associated genes.

Metaprofiles of KAKU4 binding showed a sharp, symmetric enrichment centered on gene-proximal loci in seedlings, with the strongest signal over noncoding RNA genes (tRNAs, lncRNAs) (Fig. 3D). TE sequences were consistently depleted from KAKU4 peaks. Although overall signal amplitude declined at 120 DAS, the rank order of chromatin preference remained (tRNA/lncRNA ≫ TEs). Thus, while KAKU4’s chromatin associations contracts with age, it remains preferentially associate with euchromatic regions and excluded from TE-rich heterochromatic regions. Supporting this pattern, 10 DAS peaks were enriched in AT-rich open chromatin regions (Supplementary Fig. 10C). Functional annotation of age-biased peaks highlight GO enrichment of 10 DAS KAKU4 peaks in hormone and stress-responsive pathways, including ethylene and jasmonate signaling, fatty-acid responses, wounding, and other signal-transduction categories (Fig. 3E). Peaks detected in senescence tissues at 120 DAS retained partial overlap with these categories, though with reduced effect sizes, indicating persistence of stress-related associations despite global contraction of KAKU4 occupancy.

### KAKU4 anchors to stress- and senescence-associated transcription factor networks

Integration of RNA-seq data with KAKU4-eYFP ChIP-seq profiles revealed that many Transcription Factor (TF) genes—particularly members of *WRKY*, *MYB* and *ERF* families known to regulate *Arabidopsis* leaf-senescence^31,32^—were misregulated in NE mutants, with the strongest deregulation observed in *kaku4* (Fig. 3F). Some of these loci also physically interact with KAKU4 in wild-type plants (Fig. 3F; right columns). Notably, *WRKY* genes that were upregulated in *kaku4* displayed significant KAKU4 binding in 10 DAS seedlings with 1.5-2.0-fold enrichment across *WRKY7/11/17/28/69* (Fig. 3F-H). The age-dependent contraction of promoter-proximal KAKU4 binding coincides with the family-wide transcriptional sensitivity observed in *kaku4*, although the data do not by themselves distinguish whether KAKU4 acts as a repressive or permissive scaffold at these loci (Fig. 3F-H).

### KAKU4 occupancy concentrates at discrete euchromatic chromatin states and is depleted from constitutive heterochromatin

Projection of the KAKU4-eYFP ChIP-seq peaks onto the 36-state *Arabidopsis* chromatin segmentation model (PCSD)^33^ reveals a pronounced compartment bias. Both the frequency of KAKU4 peaks and their normalized signal intensity were strongly enriched in open, regulatory euchromatin, while being reciprocally depleted from constitutive heterochromatin (Fig. 4A, Supplementary Fig. 10D). KAKU4 binding was maximally concentrated in chromatin states (CS) 16-29, annotated as highly accessible, promoter- and enhancer-like states, consistent with its euchromatic distribution (Fig. 3, 4B). In senescent tissues (120 DAS), both peak frequency and signal intensity across these states were drastically reduced. This euchromatic binding profile of KAKU4 contrasts sharply with metazoan lamina-associated proteins, which predominantly localize to silent, heterochromatic lamina-associated domains (LADs).^34,35^ Thus, in plants, KAKU4 defines a distinct architectural paradigm in which NE components preferentially associate with active, euchromatic regions rather than repressive heterochromatin domains.

**Figure 4.**
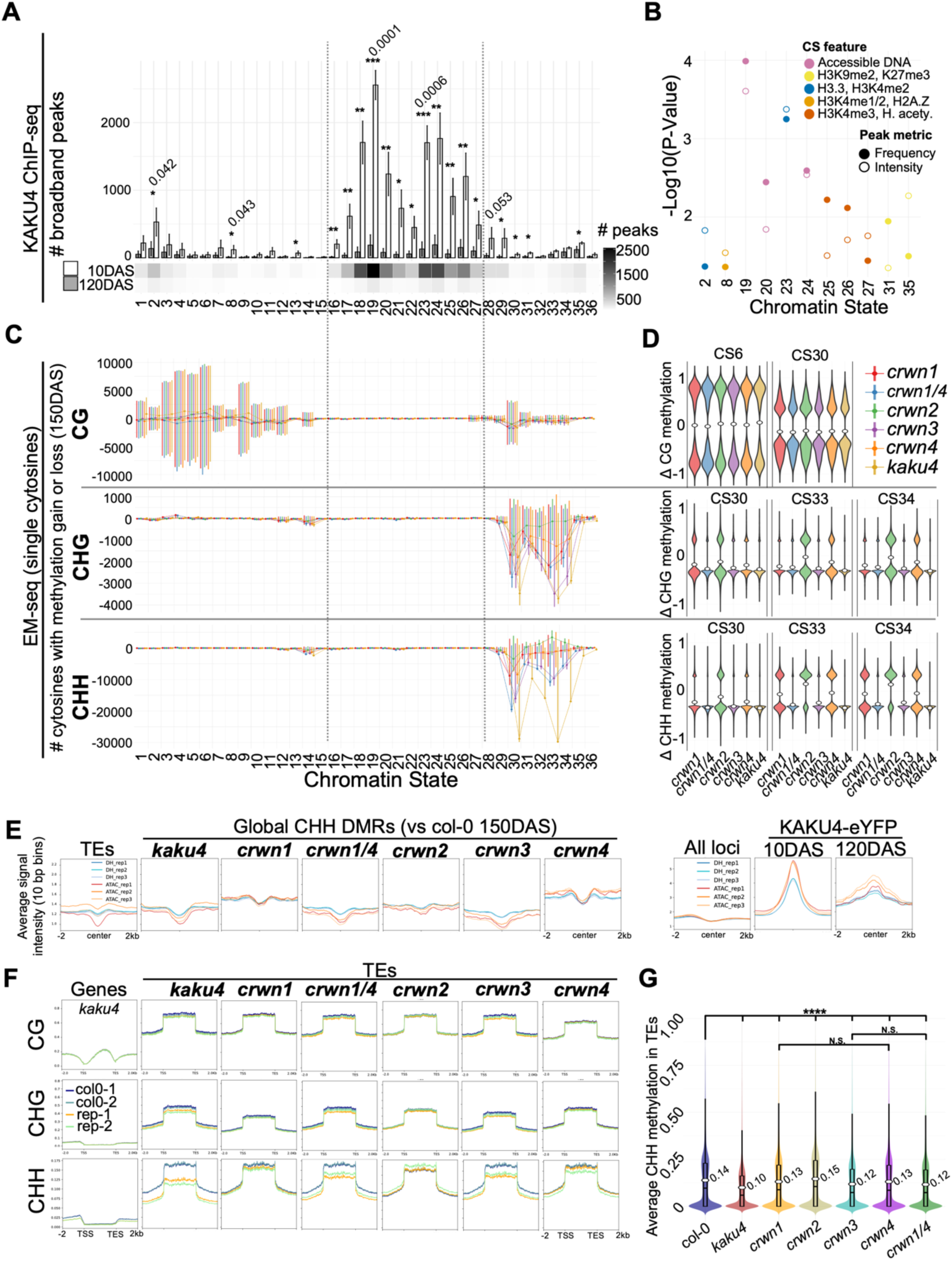
DNA methylation loss at discrete chromatin states defines an epigenomic signature of nuclear-envelope dysfunction that contrasts with KAKU4 occupancy. **(A)** Distribution of KAKU4–eYFP ChIP-seq peaks across 36 *Arabidopsis* chromatin states (PCSD model). Bars represent average peak frequency per state with standard deviation for n = 3; grey = 120 DAS, white = 10 DAS, grayscale indicates frequency of peaks. **(B)** Chromatin states with most significant change during aging for peak frequency and intensity (Sup. Fig. 10D), calculated as -log₁₀(p-value). Color labels are for main features of relevant chromatin state. **(C)** Chromatin-state-resolved methylation drift highlights heterochromatin-targeted CHH and CHG loss in nuclear-envelope mutants. Bars indicate the number of significant single differentially methylated cytosines (dots are the number of gain -loss DMCs, mutant vs Col-0 150 DAS) at 36 Chromatin States. Significant DMCs have minimum 4 reads per cytosine, 30 % methylation variance, and *p*-value threshold of 0.05, in each context and chromatin state, for each mutant line. **(D)** Violin plots for selected chromatin states from Sup. Fig. 11, showing distribution of per-cytosine DNA methylation differences (Δ methylation = mutant -WT proportion) stratified by cytosine context: CG (top), CHG (middle), and CHH (bottom). **(E)** Metaprofiles of ATAC-seq and DNase I hypersensitive assay datasets from PCSD intercepted to mutants CHH DMRs at 150 DAS (left) and KAKU4 peaks (right)*. S*ignal centered on high-confidence KAKU4-YFP peaks and DMRs (10 bp window, ±2 kb from summit). **(F)** Cytosine methylation profiles across transposable elements in nuclear-envelope mutants for two biological replicates. Line plots depict CG, CHG and CHH methylation metaprofiles over full-length annotated TEs, and genes for *kaku4* (10 bp binning, ±2 kb flanking regions). Annotations are aligned by transcription start site and transcription end site, with flanking upstream/downstream regions included for context. **(G)** Violin and embedded box plots of CHH methylation levels across TEs in Col-0 and nuclear-envelop mutants. Each data point is the mean CHH methylation level of single TE. Violin width reflects the density of TE-level methylation values; boxplot shows the median (bar), interquartile range (box), and whiskers (1.5× IQR). White dots denote group means. Pairwise Brunner–Munzel tests with Benjamini–Hochberg correction (**** FDR ≤ 0.0001, * ≤ 0.05, N.S. > 0.05).

### Nuclear-envelope mutants exhibit CHH methylation erosion in transposable elements, decoupled from KAKU4 occupancy

To determine whether NE dysfunction impacts DNA methylation stability in specific chromatin domains, we performed whole-genome methylome analysis of wild-type (Col-0), *crwn*, and *kaku4* mutants at 150 DAS. Differentially methylated cytosines (DMCs) were identified for each sequence context (CG, CHG, and CHH) and intersected with the 36 CS model. For each state, we computed the net DMC balance (number of gains minus losses), where each gain or loss represented a robust ≥30% change in methylation relative to wild-type at 150 DAS. This metric reflects cytosine-level changes capturing coordinated alterations within a given chromatin state. In parallel, per-site Δ-methylation distributions (mutant -WT) were calculated to assess the directionality and dispersion of methylation changes at single-nucleotide resolution. Across all NE mutants (*crwn1-4*, *crwn1 crwn4*, *kaku4*), a spatially restricted CHH methylation loss was pronounced (Fig. 4C, D, bottom). The most negative DMC balances occurred in H3K9me2-enriched, TE-dense chromatin states, particularly CS30, CS33, and CS34, corresponding to pericentromeric heterochromatin (Fig. 4D; Supplementary Fig. 11). In these states, tens of thousands of CHH sites were eroded, most severely in *kaku4*. By contrast, euchromatic states (for example, CS3-6) exhibited near zero net balances and tightly centered Δ-methylation distributions, indicating little to no systematic change. The broad skew of CHH Δ-methylation distributions in TE-rich states reflects a widespread loss of methylation, consistent with impaired maintenance across entire domains rather than local demethylation. These same heterochromatic states also showed substantial CHG loss (Fig. 4C, D, middle).

In the CG context, methylation loss was again enriched in TE-rich heterochromatin, but with smaller magnitude than CHG and CHH (Fig. 4C, D, top). Several euchromatic states displayed CG methylation changes in both directions, yet overall Δ-methylation distributions remained centered near zero across all mutants. The relatively stable CG gain/loss ratio across the CSs are consistent with their previously described “clocklike” epimutation behavior.^36^

We next compared the spatial relationships between open-chromatin accessibility and methylation erosion. Metaprofiles of public DNase-I hypersensitivity and ATAC-seq datasets centered on mutant-specific CHH DMRs (Δm ≥ 0.10) revealed strong depletion of accessibility signals at DMR midpoints in all NE mutants (Fig4E, left), indicating that CHH methylation erosion predominantly occurred within inaccessible, heterochromatic regions. Conversely, when the same accessibility datasets were centered on KAKU4 peaks, we observed sharp central enrichment of the open chromatin signatures (Fig. 4E, right). These reciprocal profiles delineate two distinct spatial domains: the “geography of methylation failure” in closed, TE-rich heterochromatin, and the “geography of KAKU4 occupancy” in open, gene-regulatory chromatin.

To define the most affected compartments, we generated genome-wide methylation metaprofiles across TEs. Relative to wild-type plants (150 DAS), all NE mutants showed reduced CHH methylation across TE bodies, with the strongest reduction in *kaku4* (Fig. 4F, bottom, 4G). CHG methylation displayed a parallel but less pronounced decline (middle), whereas CG methylation remained largely stable (top). Gene-centered methylation profiles were unchanged across all contexts, confirming that methylation erosion is confined to TE-dense heterochromatin and spares gene-rich euchromatic regions.

Collectively, these findings demonstrate that NE mutants undergo pronounced CHH methylation erosion specifically within heterochromatic, TE-rich domains where KAKU4 is normally absent. The clear spatial dissociation between KAKU4 occupancy and methylation loss suggests an indirect mechanism—where perturbation of nuclear architecture disrupts perinuclear silencing environments, compromising the recruitment or stability of pathways required for heterochromatin maintenance.

### Nuclear-envelope factors are required for age-associated CHH hypermethylation of heterochromatic TEs

To elucidate the chronological origin of heterochromatic CHH methylation loss in NE mutants, we compared DNA methylation profiles of young (12 days after sowing, DAS) and aged (150 DAS) plants in wild-type (*Col-0*) and NE mutants. Strikingly, the *kaku4* mutant at 12 DAS exhibited global methylation patterns nearly identical to wild-type, indicating that methylome establishment during early development is largely unaffected (Fig. 5A,B). In contrast, CHH methylation in wild type increased significantly from 12 DAS to 150 DAS (*p* = 0.004), revealing an age-associated CHH hypermethylation process during vegetative aging (Fig. 5A,B). By 150 DAS, CHH methylation levels diverged markedly among genotypes: *kaku4* displayed the lowest global CHH methylation, comparable to that of 12 DAS wild-type, while *crwn* mutants showed intermediate reductions (Fig. 5A,B). Grouping genotypes by aging-phenotype severity—Young (12 DAS *Col-0* and *kaku4*), Mild (*crwn1*, *crwn2*, *crwn4*), Severe (*kaku4*, *crwn3*, *crwn1/4*), and Control (150 DAS Col-0)—demonstrated that CHH methylation levels scale inversely with aging severity: mutants with pronounced premature-aging phenotypes retained methylation levels indistinguishable from the Young group (Fig. 5B).

**Figure 5.**
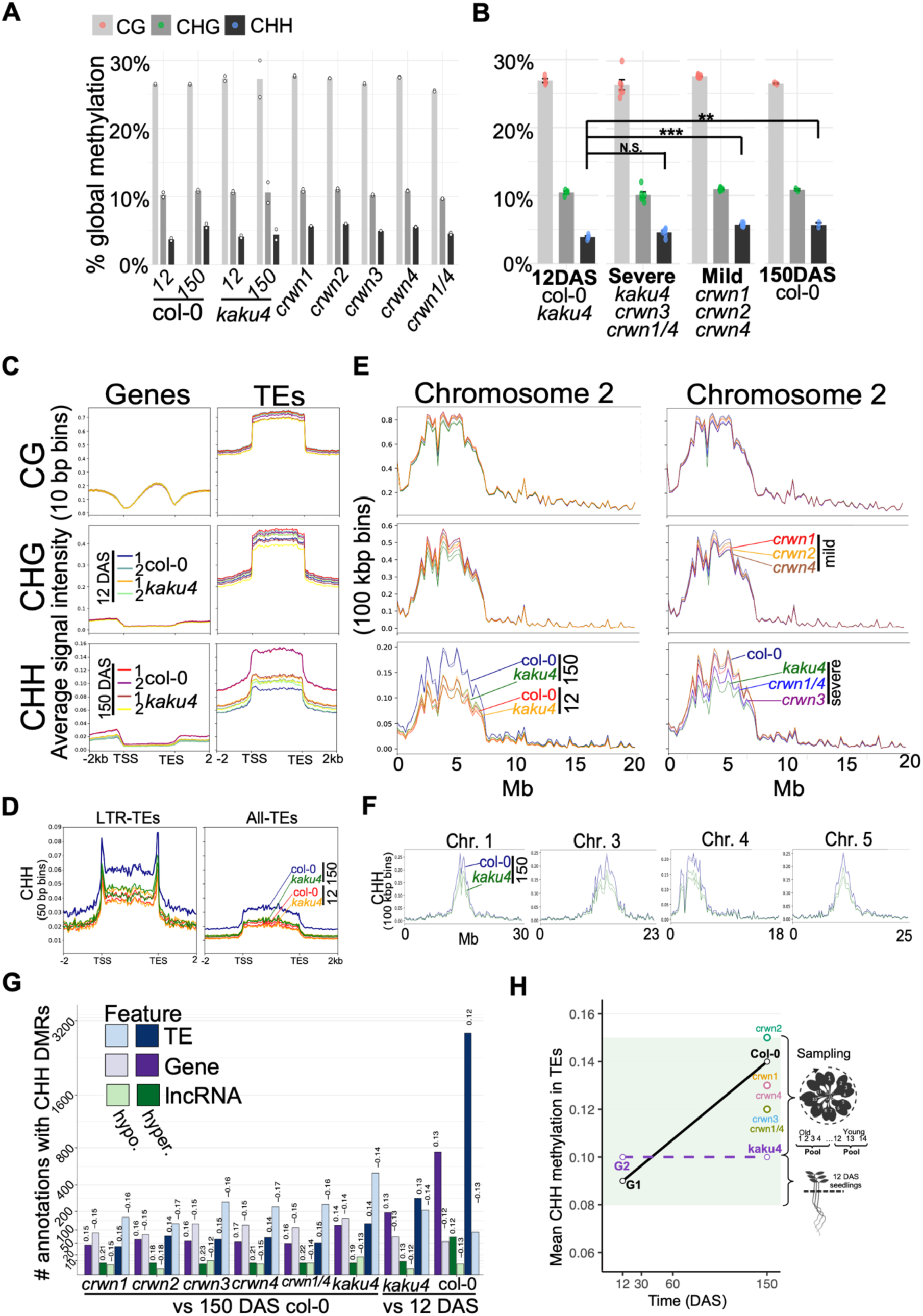
Nuclear envelope dysfunction abolishes age-linked CHH methylation gain in heterochromatic TEs. **(A)** Global methylation of nuclear-envelope mutants at 150 DAS, Col-0 at 12 and 150 DAS, and *kaku4* second generation (G2) at 12 DAS, stratified by cytosine context. **(B)** Global methylation stratified by phenotype severity class, with pairwise comparisons for CG (Young vs Mild *p* = 0.031 *), CHG (Young vs Severe *p* = 0.620; Young vs Mild *p* = 0.028 *), and CHH (Young vs Severe *p* = 0.051, N.S.; Young vs Mild *p* = 5 × 10⁻⁶ ***; Young vs Col-0 15o DAS, *p* = 0.004 **). **(C)** Genome-wide cytosine methylation profiles (CG, CHG, CHH) over TEs (±2 kb flanking), shown two replicates per condition of Col-0 and *kaku4* at 12 and 150 DAS. Line plots show methylation in Col-0 (12 DAS and 150 DAS) and *kaku4*, revealing age- and mutation-associated CHH loss over TE bodies. **(D)** CHH methylation metaprofiles over long heterochromatic LTR retrotransposons (LTR-TEs) and all transposons (All-TEs). Two replicates per color shown. **(E)** Chromosome-scale CG, CHG and CHH methylation along chromosome 2 arms (binned in 100 kb windows. Left, Col-0 and *kaku4* (12 DAS, 150 DAS) with two replicates. Right, Mild and Severe categories at 150 DAS, shown one replicate per genotype for simplicity. **(F)** CHH methylation profile along chromosomes of *A. thaliana*. Aged Col-0 (blue) and *kaku4* (green) at 150 DAS. Data of two replicates are shown. Expanded profiles for all genotypes are in Sup. Fig. 13-14. **(G)** Annotation-resolved counts of CHH DMRs overlapping TEs, protein-coding genes, and lncRNAs. DMRs were identified with thresholds (CG Δm ≥ 0.25; CHG Δm ≥ 0.10; CHH Δm ≥ 0.10; FDR q < 0.05). Light color bars denote hypomethylated and dark bars hypermethylated DMRs. Numbers above each bar indicate the mean Δm for each group. **(H)** Mean CHH methylation across full-length annotated TEs is plotted against time. Values are derived from per-TE averages of cytosine-level EM-seq data and represent replicate-merged means (n = 2 per condition). Col-0 shows an age-linked increase in TE-CHH between 12 and 150 DAS, whereas *kaku4* and other NE mutants fail to acquire the gain. *kaku4* next generation (G2) seedlings remain low, and by 150 DAS *kaku4* plants retain depressed TE-CHH relative to Col-0. Additional pointss (*crwn1*, *crwn2*, *crwn3*, *crwn4*, *crwn1/4*) show intermediate values.

Metaprofiles across full-length annotated transposable elements (TEs; ±2 kb) revealed clear age- and genotype-dependent patterns (Fig. 5C). In wild-type, CHH methylation increased with age over TE bodies. This reproducible feature across replicates suggests progressive reinforcement of CHH-mediated TE silencing during aging. CHH methylation over TE bodies is maintained by the H3K9me2-dependent pathway, whereas methylation at TE edges and short TEs is guided by the RdDM pathway.^8^ The failure of CHH methylation gain in NE mutants occurred at TE bodies, whereas border-proximal CHH changes are modest edges (Fig. 5D), consistent with impaired maintenance within H3K9me2-rich interiors, while boundary-proximal sites show comparatively minor changes.

Chromosome-scale methylation maps extended this pattern, showing age-linked CHH gains concentrated in pericentromeric heterochromatin of wild-type plants (Fig. 5E-F; Supplementary Figs. 13–14). In contrast, NE mutants failed to exhibit these age-dependent increases: CHH methylation across pericentromeric regions remained depressed at 150 DAS, with the strongest reductions in *kaku4*, *crwn3*, and *crwn1 crwn4*, and intermediate losses in *crwn1*, *crwn2*, and *crwn4* (Fig. 5C–F).

DMR analysis further revealed a TE-centric signature of aging and its disruption in NE mutants (Fig. 5G; Supplementary Fig. 12). In longitudinal comparisons of wild type and *kaku4* (12 → 150 DAS), most CHH DMRs overlapping TEs gained methylation, whereas hypomethylated DMRs were comparatively rare, and only a small fraction intersected protein-coding genes. In all NE mutants, this balance was inverted—CHH methylation losses predominated across TEs, with *kaku4* and *crwn1 crwn4* showing the highest numbers of hypomethylated DMRs (Fig. 5G).

A synthesis of CHH methylation trajectories (Fig. 5H) highlights that in wild-type rosettes, TE-linked CHH methylation increases steadily from juvenile seedlings to 150 DAS, reflecting a global age-associated gain. This rise is entirely absent in NE mutants: *kaku4* seedlings at 12 DAS, begin with low CHH levels and remain low throughout life, and mature *kaku4* rosettes at 150 DAS retain depressed CHH methylation despite advanced phenotypic age. *CRWN* paralogs trace intermediate trajectories—*crwn2* approaching wild type, while *crwn1 crwn4* lags—consistent with a graded dependence of CHH accumulation on perinuclear architecture.

Together, these findings demonstrate that CHH methylation gains during aging are not autonomously encoded in the methylation machinery but require functional nuclear-envelope components. The direction of methylome drift—an upward reinforcement of CHH methylation in heterochromatic TEs—is conditional on KAKU4- and CRWN-dependent nuclear organization, linking perinuclear organization to heterochromatin maintenance.

### Nuclear-envelope factors modulate age-associated transitions of chromatin states

To further explore how KAKU4 influences euchromatic regulation, we integrated enzymatic methyl-seq (EM-seq), KAKU4 ChIP-seq, and public chromatin datasets from the PCSD.^33^ We found that age-acquired (Col-0, 12 → 150 DAS) and mutant-acquired (*kaku4 vs.* Col-0, 150 DAS) DMRs were highly enriched near binding sites of RdDM components, including Pol IV, Pol V (NRPE1), MORC1, and JMJ14 (Fig. 6A), which typically target short euchromatic TEs and the edges of long TEs.^8^

**Figure 6.**
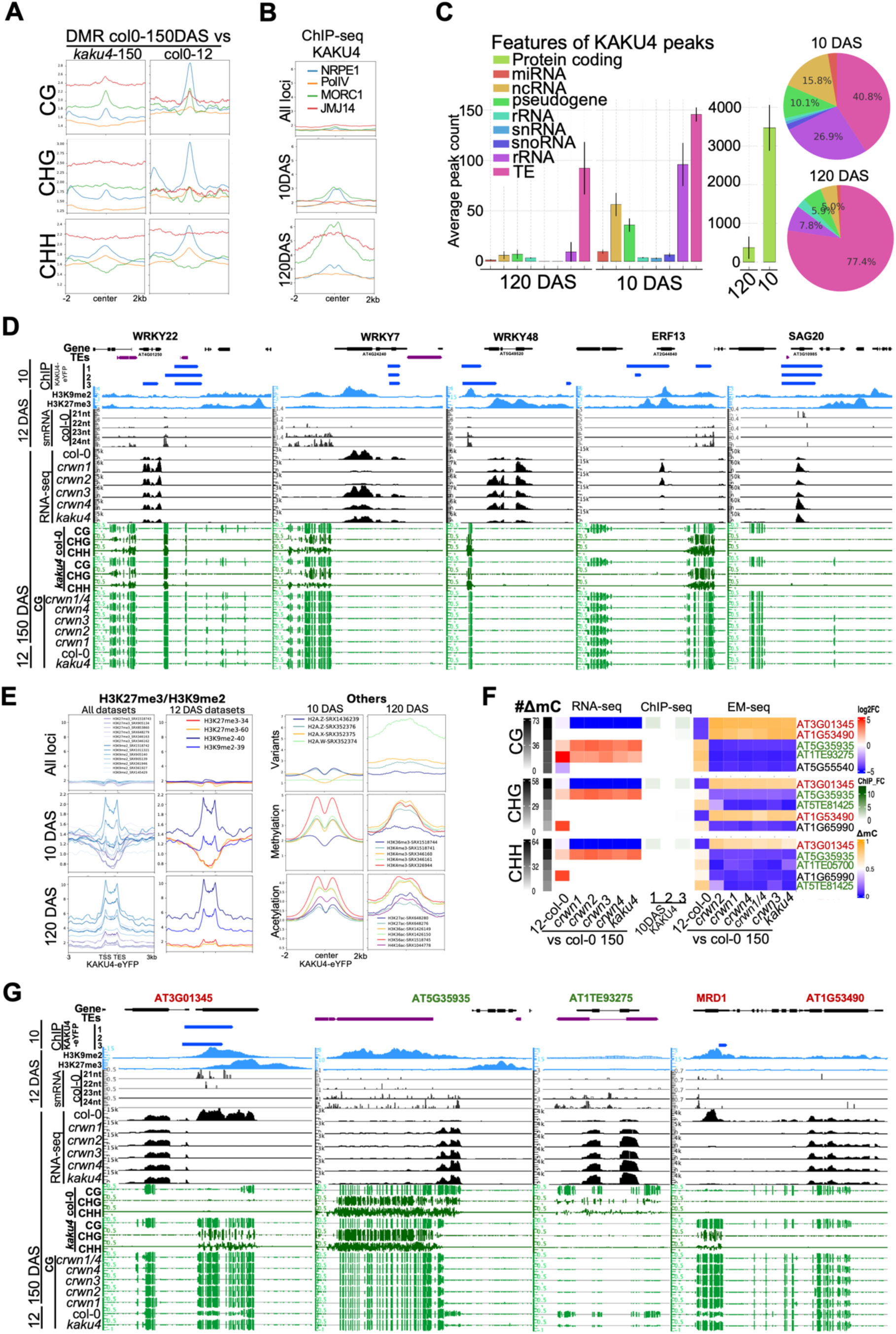
Age- and mutation-associated DMRs overlap RdDM targets linking RNA biosynthesis to mCHH heterochromatin collapse. **(A)** Global CG, CHG and CHH DMRs metaprofiles (±2 kb; CG Δm ≥ 0.25; CHG Δm ≥ 0.10; CHH Δm ≥ 0.10; FDR q < 0.05) centered on public ChIP-seq-defined binding sites (PCSD) for PolIV and RdDM components NRPE1, MORC1, and JMJ14. **(B)** 10- and 120-DAS KAKU4-eYFP metaprofiles (±2 kb) centered on public ChIP-seq-defined binding sites for PolIV and RdDM components NRPE1, MORC1, and JMJ14. **(C)** Classification of KAKU4 peaks by feature biotype (e.g., pseudogene, TE, rRNA, snRNA, snoRNA). Left, barplot showing average peak count of 120 vs 10 DAS; right, pie plot showing percentage of annotation (excluding protein-coding) and change in 120 and 10 DAS. **(D)** Genome-browser views at stress/senescence-associated transcription factors enriched for KAKU4 binding. Top: Shown are WRKY22 (AT4G01250), WRKY7 (AT4G24240), WRKY48 (AT5G49520, label in panel), ERF13 (AT2G44840), and SAG20 (AT3G10985), which were among the loci with strongest KAKU4 ChIP enrichment in Fig. 3. Second: Public ChIP-seq datasets for H3K9me2 (10–12 DAS leaves; SRX905139–40) and H3K27me3 (10–12 DAS leaves and seedlings; SRX905134, SRX853860). Third: smRNA-seq data (Col-0, 5-week leaves) with two replicates merged per read length (21–24 nt). Tracks represent combined signal from both strands. Fourth: RNA-seq coverage (black) from Col-0 and NE mutants (12 and 150 DAS). Bottom: cytosine methylation (green), shown as CG for all genotypes and all sequence contexts for Col-0/*kaku4*. Gene models are color-coded by feature type where red annotations contain ncRNAs and green are TE targets of RdDM. **(E)** Left, metaprofiles with scale regions on KAKU4-eYFP peaks (±3 kb, 1 kb TSS/TTS) of public ChIP-seq datasets for all H3K9me2 and H3K27me3 (left) and selected datasets for 10-12 DAS tissues, as described in Panel D (right). Right, metaprofiles (±2 kb) of public ChIP-seq datasets for histone variant, methylation and acetylation marks of young *Arabidopsis* leaves and seedlings (12 days–3 weeks old). **(F)** Multi-modal annotation sorted by highest frequency of cytosines (#ΔmC) in DMCs-enriched features across mutants. Left: log₂ fold-change from RNA-seq (Col-0 12 vs 150 DAS, and mutant vs Col-0 150 DAS). Center: KAKU4 ChIP-seq fold enrichment signal (ChIP/Input). Right: Average methylation in annotation across all CG, CHG, and CHH DMCs methylation profile clustered by Euclidean distance. **(G)** Genome browser tracks for representative loci identified in panel F: *AT3G01345*, *AT5G35935*, *AT1TE93275*, and *AT1G53490*. Top: KAKU4-eYFP ChIP-seq (10 DAS seedlings, broadband peaks). Tracks are explained in Fig. 6D.

Meta-analysis of Pol IV/V signals showed a characteristic twin-summit enrichment flanking KAKU4 peaks at both 10 DAS and 120 DAS (Fig. 6B), indicating that KAKU4-bound sites lie adjacent to or border RdDM targets. Annotation of KAKU4 peaks confirmed that most binding regions correspond to protein-coding genes, consistent with its promoter/TSS-enriched distribution in euchromatin (Figs. 3, 6C), while a minor subset of peaks overlapped with TEs, likely representing short TE fragments embedded in gene-rich regions. Notably, several stress- and senescence-related transcription factor loci—*WRKY22*, *WRKY7*, *WRKY48*, *ERF13*, and *SAG20*—displayed strong KAKU4 binding in seedlings, were associated with small-RNA clusters of heterochromatic character and were transcriptionally deregulated in *kaku4* at 150 DAS (Fig. 6D). These features point to KAKU4-bound euchromatic regions as potential “anchor points” maintaining the spatial segregation of chromatin states and gene regulation.

Integration with public histone-modification datasets of young tissues (seedlings and leaves, 12 days to 3 week-old) centered on KAKU4 peaks revealed a promoter-like configuration, i.e. H2A.Z nucleosomes, sharp H3K4me3, K36me3, and H3 acetylation enrichments, in flanking regions of KAKU4 binding sites (Fig. 6E, right). H3K9me2 displayed bilateral enrichment flanking KAKU4 sites, whereas H3K27me3 was largely depleted from these regions in young tissues (Fig. 6E, left). This places KAKU4 within open, promoter-proximal neighborhoods bordering repressive chromatin domains, consistent with H3K9me2 depletion at lamina-associated domains in *kaku4*,^37^ a nucleosome-proximal KAKU4/PNET2/CRWN scaffold at the nuclear lamina,^26^ and adjacent to RdDM targets inferred from twin-summit profiles of RdDM factors (Fig. 6A,B).

Loci showing the strongest methylation and expression changes in NE mutants exhibited dynamic chromatin-state transitions with age in wild-type plants (Fig. 6F,G). For example, *AT3G01345* and *AT1G53490–MRD1(AT1G53480)*, located near KAKU4 peaks, progressively lost DNA methylation during aging, whereas euchromatic TEs such as *AT5G35935* and *AT1TE93275*—canonical RdDM/Pol V targets whose silencing is compromised in RdDM mutants^38,39^—showed age-associated methylation gains across TE bodies in wild-type, similar to the age-associated DNA methylation accumulations in heterochromatic TEs (Figs. 4, 5). These age-linked chromatin-state transitions were abolished in all NE mutants (Fig. 6F,G), suggesting that nuclear-envelope proteins regulate age-linked chromatin dynamics either directly or indirectly through perinuclear interactions.

Taken together, these findings indicate that NE proteins bound to euchromatic regions act not only as structural anchors but also as regulators of local chromatin environments during aging. Through perinuclear interactions, NE proteins likely safeguard chromatin-state boundaries and mediate age-associated transitions between euchromatic and repressive domains around euchromatic TEs.

## Discussion

Our study identifies nuclear-envelope (NE) integrity as an upstream, rate-limiting determinant of epigenetic fidelity in *Arabidopsis thaliana*. Under SD conditions that separate chronological time from developmental phase, disruption of *KAKU4* and *CRWN* paralogs reveals a photoperiod-sensitive, progeroid-like trajectory—premature leaf senescence, loss of meristem competence, and accelerated transcriptional drift. These phenotypes indicate that NE dysfunction exposes a latent, time-dependent decline rather than merely advancing developmental transitions. The parallels with animal laminopathies—where lamin mutations and senescence-associated LMNB1 loss remodel LADs and late-replicating heterochromatin^9,35^—underscore a conserved principle: peripheral nuclear architecture constrains the trajectory of functional decline across eukaryotes.^1,3^

Transcriptome analyses position NE mutants along the wild-type aging axis, reflecting acceleration of intrinsic stress and aging programs. Two coordinated expression arms emerge: constitutive activation of WRKY/NAC/ERF regulons that govern defense and senescence, and suppression of nucleolar and RNA-biosynthetic modules that sustain growth. The quantitative hierarchy (*kaku4* ≫ *crwn1 crwn4* ≫ single *crwns*) mirrors phenotypic severity, identifying KAKU4 as the most sensitive node coupling RNA metabolism to chromatin homeostasis. Consistent with a lamina-based regulatory interface, CRWNs and their partners interact with Polycomb-group (PcG) and repressive complexes,^12,13,40^ embedding transcriptional restraint within the mechanical framework of the nuclear periphery. The plant lamina also dynamically reorganizes genome topology under stress^12^ and can nucleate phase-separated responses at the nuclear rim.^41^

Mechanistically, KAKU4 ChIP-seq data delineates a promoter- and TSS-enriched occupancy atlas with minimal TE overlap (Figs. 4, 6). With age, this landscape contracts approximately eightfold and shifts from sharp promoter anchors toward broader genic contacts (Figs. 4, 6). Such redistribution and contraction mirror promoter detachment from lamina contacts in mammalian senescence^10,42,43^ and aligns with studies showing position-dependent gene regulation via perinuclear tethering in plants.^44,45^ Persistent KAKU4 binding at tRNA and ncRNA hotspots—regions exhibiting boundary or insulator-like behavior in plants and animals^46,47^—suggests that KAKU4 stabilizes promoter insulation and gates stimulus-responsive loci. Thus, the plant lamina functions less as a static heterochromatin tether and more as a dynamic scaffold for euchromatic regulation.

In contrast, methylome drift in NE mutants is compartmentalized and architecture-dependent: CHH and CHG methylation loss concentrates within H3K9me2-dense pericentromeric heterochromatin. Critically, these losses occur outside KAKU4 occupancy zones, excluding a direct tether model.^8,38,39,48,49^ This spatial decoupling parallels observations in mammalian laminopathies, where lamina-associated hypomethylation arises distal to transcription factor binding sites.^9,10,35,50^ Across kingdoms, NE integrity thus maintains heterochromatin stability indirectly—by preserving the structural and biochemical milieu required for methylation maintenance rather than through direct chromatin contact.

In summary, when perinuclear scaffolds remain intact, heterochromatin retains methylation and transcriptional stability; when they fail, reinforcement of developmental silencing collapses and epigenetic drift accelerates. Despite deep evolutionary rewiring of lamina composition between kingdoms,^1,4,13^ the functional imperative to stabilize repressive domains remains conserved.^9,12,35^ Thus, nuclear-envelope integrity emerges as a conserved regulator of chromatin stability and epigenetic aging, and progeroid NE mutants in *Arabidopsis* provide a tractable framework to uncover how nuclear architecture influences the rate and reversibility of aging.

## Acknowledgments

We thank Drs. Tamura and Nishimura for providing seeds of KAKU4-eYFP line and Drs. Sakamoto and Matsunaga for seeds of *crwn* mutants. We thank the OIST SQC for RNA-seq, ChIP-seq and EM-seq sequencing services.

## Author contributions

Conceptualization: OJ. Methodology: OJ & HS. Investigation: OJ. Visualization: OJ. Funding acquisition: HS. Supervision: HS. Writing: OJ & HS.

## Data and materials availability

All high-throughput sequencing data generated in this study have been submitted to ArrayExpress (EMBL–EBI) and will be made publicly available upon request.

RNA-seq data are under accession E-MTAB-16251

Whole-genome methylome (EM-seq) data under accession E-MTAB-16263

ChIP–seq KAKU4-eYFP dataset under accession E-MTAB-16266. All *Arabidopsis thaliana* lines used in this work are in the Col-0 background and are available from public stock centers or from the corresponding author upon reasonable request.

## Supplementary Materials

Materials and Methods

Sup. Figs. 1 to 14

Tables 1 to 4

## Materials and Methods

### Plant material and growth

All experiments used *Arabidopsis thaliana* Columbia-0 (Col-0). T-DNA lines. CRWN: *crwn1* (AT1G67230; SALK_016800), *crwn2* (AT1G13220; SALK_076653), *crwn3* (AT1G68790; SALK_099283), *crwn4* (AT5G65770; SALK_079288); alternative alleles *crwn3-b* (SALK_134237), *crwn4-b* (SALK_097845).KAKU4 (AT4G31430): SALK_076754 (principal), SALK_012298, SALK_082152, SAIL_771_E09. ProKAKU4:KAKU4-eYFP line complemented *kaku4-3* was kindly provided with Drs. Tamura and Nishimura. AtMAN1 (AT5G46560): SALK_024888 (CS887097), SAIL_392_H12 (CS818091). Regulatory/epigenetic mutants: *ATL31* (AT5G27420; SALK_002809), *ELF3* (AT2G25930; SALK_202881), *ANAC017* (AT1G34190; SALK_025104), *SSPP/APD7* (AT5G02760; SALK_099356), *bHLH03* (AT4G16430; SALK_050954), *ORE15* (AT1G31040; SALK_144267). Flowering regulators: *FUL* (*ful-a*, SALK_096481; *ful-b*, SAIL_1291_B09), *SOC1* (*soc1-a*, SALK_138131; *soc1-b*, SALK_006054).

Of the five independent *kaku4* T-DNA alleles (SALK_076754/*kaku4-2*, SALK_010298/*kaku4-3*, SAIL_771_E09/*kaku4-4*, SALK_082152, SALK_011890), SALK_076754/*kaku4-2* and SAIL_771_E09/*kaku4-4* showed the accelerated senescence phenotypes, including premature rosette collapse and bolting arrest under SD at ∼130 DAS (Sup. Fig. 3C), and SALK_076754/*kaku4-2* was used for majority of the analysis in this study. Seeds for *crwn1–4* and double homozygotes (*crwn1 crwn2*, *crwn1 crwn4*, *crwn2 crwn3*) were kindly gifted from Drs. Sakamoto and Matsunaga; other seeds were obtained from ABRC/NASC.

### Growth conditions

Seeds were surface-sterilized (15% commercial bleach; ∼0.8% NaClO, 10 min), rinsed 5×, stratified (4 °C, dark, 48 h), and sown 3–5 seeds/pot. Plants were grown in controlled chambers (Biotron LPH-410SP; NK System) at 22 °C and 65% RH under long day (LD) 16 h light/8 h dark (∼200 µmol m⁻² s⁻¹) or short day (SD) 8 h light/16 h dark (∼120 µmol m⁻² s⁻¹).

### Lifespan phenotyping

Per genotype, 9 plants (3 plants/pot × 3 pots) were scored. Phases: seedling (to cotyledon expansion), rosette development (to first visible yellowing), rosette senescence (from first yellowing to complete loss of green rosette tissue), and bolting/flowering (stem >5 mm to full extension, with +10 d buffer). Leaf senescence was scored as ≥70% yellowing/necrosis. In LD, scoring considered the oldest 5–6 leaves while 10–12 leaves in SD. Plants failing to bolt by 120 days after sowing (DAS) were recorded as vegetatively arrested; lifespans were truncated at 160 DAS. Representative plants (Col-0, *kaku4*, *crwn1–4*) were imaged at 90, 115, and 160 DAS using a standardized white-balance/exposure workflow; aborted meristems were confirmed microscopically.

### Plant sampling for omics

“Early” plant samples were collected at 12 DAS (2–4-leaf stage). “Late” multi-omics sampling was conducted at 150 DAS (SD), when phenotypes diverged maximally. From each rosette, leaves 1–4 (basal) and 11–14 (apical) were dissected and combined to represent the whole rosette. Each biological replicate pooled tissue from 3–4 plants (one pot); two biological replicates were processed per genotype, with a third pot maintained as backup. All harvests were at ZT4. Tissues were flash-frozen, and stored at −80 °C. DNA concentration was verified (Qubit dsDNA BR) prior to EM-seq.

### Replication note

Severe SD phenotypes reduced tissue yield (especially in *kaku4*), constraining omics to n=2 biological replicates. We prioritized robust, direction-consistent signals and used orthogonal agreement across RNA-seq, EM-seq, and KAKU4-eYFP ChIP-seq; internal genetic controls behaved as expected under LD/SD.

### RNA-seq

#### Library preparation and sequencing

Total RNA (100 ng) from 150 DAS SD rosettes (*crwn1–4*, *kaku4*, Col-0) and 12 DAS Col-0 was extracted (RNeasy Plant Mini, Qiagen 74904), rRNA-depleted (QIAseq FastSelect). RNA-seq libraries were prepared with NEBNext Ultra II Directional RNA (NEB E7760). rRNA-depleted RNA was fragmented at 94 °C (8 min); first-strand cDNA at 25 °C (10 min), 42 °C (50 min), 70 °C (15 min); dUTP-second strand per kit; SPRIselect 1.8× cleanup; end-repair/A-tail and adaptor ligation (NEBNext Ultra II Ligation Master Mix; USER 37 °C, 15 min); 0.9× cleanup; 12 PCR cycles (NEB Ultra II Q5). Libraries were bead-cleaned, quality-checked (Bioanalyzer HS/1000), quantified (Qubit dsDNA HS), pooled, and sequenced at OIST SQC (NovaSeq 6000, 2×150 bp). Summary of RNA-seq sequencing and alignment metrics is in **Table 1**.

#### Data Processing

polyG/polyX and 5′ quality trimming was performed by fastp v0.23.2.^51^ Alignment to TAIR10 nuclear chromosomes (Chr1–5) used HISAT2 v2.2.1 with Araport11/atRTD3-derived splice guidance (--dta --rna-strandness RF --no-mixed --max-intronlen 10000 -p 16); BAM files were sorted/indexed by samtools v1.16.^52^ Reads aligned to genes were counted with HTSeq-count v0.12.4^53^ (-r pos -s reverse -t exon -i gene_id) using Araport11 nuclear gene annotation. TE expression was quantified separately against a URGI TAIR10 TE-only GTF. Transcript-level abundance was estimated with Salmon v1.9.0^54^ quasi-mapping to an atRTD3 decoy-aware “gentrome” (--validateMappings -l A, optional --gcBias); Salmon TPMs were used for visualization only. Differential expression tests were performed by HTSeq-counts.

### KAKU4 ChIP-seq

#### Plant material

A functional ProKAKU4:KAKU4-eYFP complemented for *kaku4-3* (T3 homozygous; hygromycin-selected) and an untagged *kaku4* control (SALK_076754) plants were used. Young samples: 10-day seedlings on ½ MS under LD (12 seedlings/replicate; n=3 per condition). Old samples: 120 DAS SD rosettes, pooling leaves 1–4 from 3–4 plants/pot (n=3). Sample collections were conducted at ZT4.

#### ChIP

Per replicate, 0.5 g (10 DAS) or 1 g (120 DAS) tissue were frozen, ground in liquid nitrogen, and resuspended in 11 mL of ice-cold nuclei isolation buffer (10 mM HEPES pH 7.6, 1 M sucrose, 5 mM KCl, 5 mM MgCl₂, 5 mM EDTA) freshly supplemented with 0.78 mL of 16% formaldehyde (Pierce), 0.375 mL of 20% Triton X-100, 12.5 μL of 2-mercaptoethanol (Wako), and 125 μL Complete tablet solution (Roche, 1 tablet in 0.5 mL H₂O). Crosslinking was carried out for 15 minutes at room temperature with periodic inversion, followed by quenching with 0.85 mL of 2 M glycine for 5 minutes. Nuclei were enriched with 15% Percoll, and lysed by SDS lysis buffer (50 mM Tris-HCl pH 7.8, 1% SDS, 10 mM EDTA), and diluted by Buffer 1 (50 mM HEPES-KOH pH 7.6, 140 mM NaCl, 1% Triton X-100, 0.1% sodium deoxycholate, 1 mM EDTA). Genome DNA was sonicated (Branson Digital; 12×: 30 s on/1 min off; 50% amplitude) to 250–350 bp. Anti-GFP Immuno Precipitation (IP) was performed with Dynabeads pre-bound with anti-GFP (Cell Signaling #2555; ∼1:1600 dilution) for 1 h at 4 °C. Lysates (including the untagged control) were incubated with the Antibody-Dynabeads solution for 2 h at 4 °C with rotation. For washing and reverse crosslinking, the magnet-bound beads was washed with Buffer 1 (twice), Buffer 2 (10 mM Tris-HCl pH 8.0, 250 mM LiCl, 0.5% NP-40, 0.5% Na-deoxycholate), and TE (10 mM Tris-HCl pH 8.0, 1 mM EDTA). Precipitated chromatin was eluted in TE and RNase A, incubated at 37 °C for 30 min, followed by Proteinase K treatment at 42 °C for 1 h and reverse crosslinking at 65 °C for up to 14 h. DNA was purified using QIAquick columns and eluted in ultrapure water.

#### Libraries and sequencing

NEBNext Ultra II DNA (E7645) with NEBNext Multiplex Oligos (E6440) was used for library preparation with following conditions; bead size-selection >150 bp (peak 270–500 bp including adaptors); 8 PCR cycles. Pooled libraries were sequenced at SQC with NovaSeq 6000 (2×150 bp; ≥20 M fragments/sample). Summary of KAKU4-eYFP ChIP-seq and alignment metrics are in **Table 2**.

#### Data Processing

ChIP-seq data were processed using the nf-core/chipseq pipeline v2.0.0 (https://nf-co.re/chipseq),^55^ deployed on the OIST HPC cluster. Pipeline citations can be found at nf-core/chipseq for the following bioinformatic tools used: Raw FASTQ files were processed through standardized steps including adapter trimming with Trim Galore, quality control with FastQC, alignment, duplicate marking, filtering, normalization, peak calling, and downstream quantification. Reads were aligned to the *Arabidopsis* thaliana TAIR10 reference genome (excluding organellar chromosomes) using Bowtie2 (v2.4.5), with paired-end settings. Duplicates were marked with Picard MarkDuplicates and multiple libraries per sample were merged, followed by re-marking of duplicates. Post-alignment filtering was performed to remove reads; flagged as duplicates, secondary alignments, or multimappers; containing >4 mismatches or excessively large insert sizes (>2 kb); mapping across different chromosomes or in non-FR orientation (for PE reads). Filtered BAM files were used to generate counts-per-million (CPM)–normalized bigWig tracks with deepTools (v3.5.1) as implemented in nf-core/chipseq, and coverage metaprofiles represent the mean CPM-normalized signal per genomic bin. Strand cross-correlation analysis was performed using phantompeakqualtools to estimate ChIP enrichment quality metrics (NSC, RSC). Broad peaks were called with MACS2 (v2.2.7.1) using the following parameters; macs2 callpeak --broad --broad-cutoff 0.1 --qvalue 0.05 --format BAMPE --gsize 1.19e8. Peak calling was conducted independently per replicate. Peaks with q < 0.05 and broad-cutoff < 0.1 were retained. Replicate consistency was assessed using bedtools multiinter (v2.30.0), and consensus peaks (≥2 replicates) were used for downstream read quantification with featureCounts and differential peak analysis. Peak annotation relative to genomic features was performed using HOMER, and enrichment patterns were visualized in IGB^56^ using generated session files. All quality control summaries, including raw read quality, alignment statistics, duplication rates, enrichment profiles, and peak metrics, were aggregated using MultiQC in the nf-core/chipseq platform.

#### Feature/chromatin-state analyses

Motif enrichment analysis was performed using MEME-ChIP (v5.5.2)^57^ on 250 bp sequences extracted from the summit-centered regions of MACS2-called peaks (10 DAS samples). Background sequences were generated by GC-matched genomic sampling. Only motifs with E-values < 1e−50 were considered significantly enriched. Top motifs were visualized using default MEME Suite logos.^57^

#### Statistics

Comparisons of peak counts (Fig. 4A) and ChIP signal intensities (Supplementary Fig. 10D) between young and old tissues were performed using two-sided unpaired Student’s t-tests, with Welch’s correction applied when variance heterogeneity was detected. For chromatin state-resolved comparisons, per-state t-tests were conducted and adjusted using the Benjamini-Hochberg false discovery rate (FDR) procedure. All analyses involving multiple hypothesis testing, including GO term enrichment (Fig. 3E), were similarly corrected using the Benjamini-Hochberg method. Exact P-values and adjusted P-values (FDR) are reported in the source data files. Pearson correlation coefficients across biological replicates and ChIP assays were calculated using deeptools plotCorrelation.^58^

### Enzymatic methyl-seq (EM-seq)

#### Libraries and sequencing

Genomic DNA (200 ng) from Col-0, *crwn1–4*, *crwn1 crwn4*, and *kaku4* (12 DAS and/or 150 DAS SD) was sheared (Covaris, ∼250 bp; 55 µL, peak power 75 W, duty factor 10%, 200 cycles/burst, 160 s) and sequencing library was prepared with NEBNext EM-seq kit (NEB E7120) following manufacture’s instruction. Final library amplification was carried out with EM-seq Index Primer and NEBNext Q5U Master Mix, using the following PCR cycling conditions: 98 °C for 30 s, followed by 10 cycles (98 °C for 10 s, 60 °C for 30 s, 65 °C for 60 s), and a final extension at 65 °C for 5 min. Libraries were sequenced at OIST SQC with NovaSeq 6000 (2×150 bp). Summary of EM-seq sequencing and alignment metrics nuclear-envelope mutants in **Table 3**.

#### Processing and methylation calling

Raw EM-seq reads were processed using fastp (v0.23.2)^51^ for adapter removal and base quality filtering. Paired-end reads passing quality thresholds were retained for downstream mapping. Genome indices were prepared using bismark_genome_preparation (v0.23.1)^59^ with Bowtie2 for the *Arabidopsis thaliana* TAIR10. Cleaned reads were mapped using bismark in paired-end, non-directional mode (--non_directional, --score_min L,0,-0.6) to account for EM-seq library properties. Aligned BAMs were deduplicated using deduplicate_bismark and sorted using samtools (v1.16.1). Cytosine methylation extraction was performed with bismark_methylation_extractor using the --CX, --bedGraph, --cytosine_report, and --gzip flags. Cytosines with <3× coverage were excluded using context-specific filters, and the retained methylation calls were stratified for downstream analyses.

#### DMR detection

Single-cytosine binomial tests (CG/CHG/CHH) was used for a chloroplast-based background. BH-FDR q<0.05 sites were merged with bedtools merge function (no fixed bins) into regions per condition retaining strand/context/ratio/coverage information. Mutant–control Δm were classified as gain/loss with thresholds; CG ≥0.25 (stringent ≥0.40), CHG ≥0.10 (≥0.20), CHH ≥0.10 (≥0.20).

#### Feature/chromatin-state analyses and visualization

DMRs were overlapped with Araport11 genes, lncRNAs, TEs (≥1 bp) annotations. Chromatin states (PCSD 36-state)() were assigned to DMRs and individual cytosines. Whole-genome context-wise methylation landscapes were summarized in 100-kb windows with deepTools (computeMatrix, scale-regions)^58^ and plotted (plotProfile). Feature-centered profiles were generated for full-length TEs (body ±2 kb, 10 bp bins) and public TF ChIP sites (e.g., WRKY18, NRPE1, Pol IV, ERF115) ±2 kb downloaded from PCSD.^33^ For epigenomic overlap, DMRs and KAKU4 peaks were intersected with peak summits from ATACseq, DNase-seq, H3K9me2, H3K27me3, H3.1, downloaded from PCSD; signals were summarized with deeptools (reference-point) and RPKM-normalized. LTR-TE annotations were extracted with LTR-retriever.^60^

#### Single-cytosine mapping

Genome-wide single-cytosine methylation profiling was performed using CX reports derived from Bismark.^59^ These files pre-filtered to exclude organellar chromosomes and strand-separated. This step avoids collapsing methylation signals from Watson/Crick strands at CpG palindromes, which would otherwise inflate DMC width and cytosine counts. Replicates were merged using DMRcaller joinReplicates function.^61^ Differential methylation was computed separately for CG, CHG, and CHH contexts using DMRcaller computeDMRsReplicates with method = “bins” and test = “betareg”. Pseudocounts (pseudocountM = 1, pseudocountN = 2) were applied to stabilize methylation estimates near boundaries (0 or 1). Binning strategies for the single-cytosine mode (binSize = 1): DMCs required minCytosinesCount = 1, minGap = 1, minSize = 1, minReadsPerCytosine = 4, minProportionDifference = 0.3, and pValueThreshold = 0.05. Using R v4.2.0 packages (https://www.R-project.org/), filtered outputs were converted to Granges and intersected with a 36-state chromatin segmentation model using GenomicRanges::findOverlaps (v1.44). Functional state assignments (e.g., active promoters, Polycomb-repressed regions, TE-rich heterochromatin) were resolved by maximal segment coverage per DMC. Aggregation using dplyr::group_by yielded per-state summaries of gain/loss counts, net methylation, and average absolute change.

#### Statistics

R v4.2.0 (https://www.R-project.org/) was used for statistical tests: Brunner–Munzel tests were applied for per-site distributions; Wilcoxon rank-sum or unpaired two-sided t-tests where appropriate; BH-FDR throughout for multiple testing.

#### Reconciling *kaku4* replicate variation

At 150 DAS, *kaku4* replicates varied in global means (e.g., CG 30.8%/CHH 5.9% vs CG 24.9%/CHH 3.7%) despite comparable QC (conversion 99.5–99.9%, alignment 74–83%, depth 80–90×). Differences reflect leaf-cohort composition imposed by severe SD phenotypes while TE-centered profiles showed consistent CHH loss in both replicates. All CHH aging inferences derived from TE-resolved statistics were robust to cohort composition.

#### Small-RNA datasets

Preprocessed small-RNA genome-coverage tracks (WT Col-0 12 DAS seedlings) were obtained from the data in preprint (Shimada and Saze, 2024)^62^ mapped to TAIR10 and separated by read length (21, 22, 24 nt) and strand.

## References

1. López-Otín, C., Blasco, M. A., Partridge, L., Serrano, M. & Kroemer, G. Hallmarks of aging: An expanding universe. Cell 186, 243–278 (2023).

2. Owen, J. et al. Diversity of ageing across the tree of life. Nature 505, (2014).

3. Lemoine, M. The Evolution of the Hallmarks of Aging. Front. Genet. 12, 693071 (2021).

4. Mermet, S. et al. Evolutionarily conserved protein motifs drive interactions between the plant nucleoskeleton and nuclear pores. Plant Cell 35, 4284–4303 (2023).

5. Groves, N. R. et al. Recent advances in understanding the biological roles of the plant nuclear envelope. Nucleus 11, 330–346 (2020).

6. Suzuki, M. M. & Bird, A. DNA methylation landscapes: provocative insights from epigenomics. Nat. Rev. Genet. 9, 465–476 (2008).

7. Zhang, H., Lang, Z. & Zhu, J.-K. Dynamics and function of DNA methylation in plants. Nat. Rev. Mol. Cell Biol. 19, 489–506 (2018).

8. Law, J. A. & Jacobsen, S. E. Establishing, maintaining and modifying DNA methylation patterns in plants and animals. Nat. Rev. Genet. 11, 204–220 (2010).

9. Shah, P. P. et al. Lamin B1 depletion in senescent cells triggers large-scale changes in gene expression and the chromatin landscape. Genes Dev. 27, 1787–1799 (2013).

10. Chandra, T. et al. Global reorganization of the nuclear landscape in senescent cells. Cell Rep. 10, 471–483 (2015).

11. Zhou, W. et al. DNA methylation loss in late-replicating domains is linked to mitotic cell division. Nat. Genet. 50, 591–602 (2018).

12. Wang, N. et al. The plant nuclear lamina disassembles to regulate genome folding in stress conditions. Nat. Plants 9, 1081–1093 (2023).

13. Ciska, M., Hikida, R., Masuda, K. & Moreno Díaz de la Espina, S. Evolutionary history and structure of nuclear matrix constituent proteins, the plant analogues of lamins. J. Exp. Bot. 70, 2651–2664 (2019).

14. Hu, B. et al. Plant lamin-like proteins mediate chromatin tethering at the nuclear periphery. Genome Biol. 20, 87 (2019).

15. Goto, C., Tamura, K., Fukao, Y., Shimada, T. & Hara-Nishimura, I. The Novel Nuclear Envelope Protein KAKU4 Modulates Nuclear Morphology in Arabidopsis. Plant Cell 26, 2143–2155 (2014).

16. Goto, C., Hara-Nishimura, I. & Tamura, K. Regulation and Physiological Significance of the Nuclear Shape in Plants. Front. Plant Sci. 12, 673905 (2021).

17. Koornneef, M., Hanhart, C. J. & van der Veen, J. H. A genetic and physiological analysis of late flowering mutants in Arabidopsis thaliana. Mol. Gen. Genet. MGG 229, 57–66 (1991).

18. Nooden, L. D., Hillsberg, J. W. & Schneider, M. J. Induction of leaf senescence in Arabidopsis thaliana by long days through a light-dosage effect. Physiol. Plant. 96, 491–495 (1996).

19. Cao, Y. et al. KAKU4 regulates leaf senescence through modulation of H3K27me3 deposition in the Arabidopsis genome. BMC Plant Biol. 24, 177 (2024).

20. Aoyama, S. et al. Ubiquitin Ligase ATL31 Functions in Leaf Senescence in Response to the Balance Between Atmospheric CO2 and Nitrogen Availability in Arabidopsis. Plant Cell Physiol. 55, 293–305 (2014).

21. Anwer, M. U., Davis, A., Davis, S. J. & Quint, M. Photoperiod sensing of the circadian clock is controlled by EARLY FLOWERING 3 and GIGANTEA. Plant J. Cell Mol. Biol. 101, 1397–1410 (2020).

22. Broda, M. et al. Increased expression of ANAC017 primes for accelerated senescence. Plant Physiol. 186, 2205–2221 (2021).

23. Xiao, D. et al. SENESCENCE-SUPPRESSED PROTEIN PHOSPHATASE Directly Interacts with the Cytoplasmic Domain of SENESCENCE-ASSOCIATED RECEPTOR-LIKE KINASE and Negatively Regulates Leaf Senescence in Arabidopsis1[OPEN]. Plant Physiol. 169, 1275–1291 (2015).

24. Serrano-Bueno, G., Sánchez de Medina Hernández, V. & Valverde, F. Photoperiodic Signaling and Senescence, an Ancient Solution to a Modern Problem? Front. Plant Sci. 12, 634393 (2021).

25. Chen, J. et al. Suppressor of Overexpression of CO 1 Negatively Regulates Dark-Induced Leaf Degreening and Senescence by Directly Repressing Pheophytinase and Other Senescence-Associated Genes in Arabidopsis. Plant Physiol. 173, 1881–1891 (2017).

26. Tang, Y., Dong, Q., Wang, T., Gong, L. & Gu, Y. PNET2 is a component of the plant nuclear lamina and is required for proper genome organization and activity. Dev. Cell 57, 19–31.e6 (2022).

27. Brocklehurst, S. et al. Induction of epigenetic variation in Arabidopsis by over-expression of DNA METHYLTRANSFERASE1 (MET1). PLOS ONE 13, e0192170 (2018).

28. Havecker, E. R., Wallbridge, L. M., Fedito, P., Hardcastle, T. J. & Baulcombe, D. C. Metastable Differentially Methylated Regions within Arabidopsis Inbred Populations Are Associated with Modified Expression of Non-Coding Transcripts. PLOS ONE 7, e45242 (2012).

29. Choi, J., Strickler, S. R. & Richards, E. J. Loss of CRWN Nuclear Proteins Induces Cell Death and Salicylic Acid Defense Signaling1[OPEN]. Plant Physiol. 179, 1315–1329 (2019).

30. Guo, T. et al. Lamin-like Proteins Negatively Regulate Plant Immunity through NAC WITH TRANSMEMBRANE MOTIF1-LIKE9 and NONEXPRESSOR OF PR GENES1 in Arabidopsis thaliana. Mol. Plant 10, 1334–1348 (2017).

31. Cao, J., Liu, H., Tan, S. & Li, Z. Transcription Factors-Regulated Leaf Senescence: Current Knowledge, Challenges and Approaches. Int. J. Mol. Sci. 24, 9245 (2023).

32. Guo, Y. et al. Leaf senescence: progression, regulation, and application. Mol. Hortic. 1, 5 (2021).

33. Liu, Y. et al. PCSD: a plant chromatin state database. Nucleic Acids Res. 46, D1157–D1167 (2018).

34. Guelen, L. et al. Domain organization of human chromosomes revealed by mapping of nuclear lamina interactions. Nature 453, 948–951 (2008).

35. van Steensel, B. & Belmont, A. S. Lamina-associated domains: links with chromosome architecture, heterochromatin and gene repression. Cell 169, 780–791 (2017).

36. Yao, N. et al. An evolutionary epigenetic clock in plants. Science 381, 1440–1445 (2023).

37. Cao, Y. et al. Nuclear lamina component KAKU4 regulates chromatin states and transcriptional regulation in the Arabidopsis genome. BMC Biol. 22, 80 (2024).

38. Yokthongwattana, C. et al. MOM1 and Pol-IV/V interactions regulate the intensity and specificity of transcriptional gene silencing. EMBO J. 29, 340–351 (2010).

39. Pontier, D. et al. NERD, a plant-specific GW protein, defines an additional RNAi-dependent chromatin-based pathway in Arabidopsis. Mol. Cell 48, 121–132 (2012).

40. Yang, T. et al. PWOs repress gene transcription by regulating chromatin structures in Arabidopsis. Nucleic Acids Res. 52, 12918–12929 (2024).

41. Tang, Y. et al. Nuclear lamina phase separation orchestrates stress-induced transcriptional responses in plants. Dev. Cell S1534-5807(25)00446–0 (2025) doi:10.1016/j.devcel.2025.07.008.

42. Finlan, L. E. et al. Recruitment to the nuclear periphery can alter expression of genes in human cells. PLoS Genet. 4, e1000039 (2008).

43. Kind, J. et al. Genome-wide maps of nuclear lamina interactions in single human cells. Cell 163, 134–147 (2015).

44. Bi, X. et al. Nonrandom domain organization of the Arabidopsis genome at the nuclear periphery. Genome Res. 27, 1162–1173 (2017).

45. Sakamoto, Y. et al. Subnuclear gene positioning through lamina association affects copper tolerance. Nat. Commun. 11, 5914 (2020).

46. Lucero, L., Fonouni-Farde, C., Crespi, M. & Ariel, F. Long noncoding RNAs shape transcription in plants. Transcription 11, 160–171 (2020).

47. Mattick, J. S. et al. Long non-coding RNAs: definitions, functions, challenges and recommendations. Nat. Rev. Mol. Cell Biol. 24, 430–447 (2023).

48. Matzke, M. A. & Mosher, R. A. RNA-directed DNA methylation: an epigenetic pathway of increasing complexity. Nat. Rev. Genet. 15, 394–408 (2014).

49. Wang, Z. & Baulcombe, D. C. Transposon age and non-CG methylation. Nat. Commun. 11, 1221 (2020).

50. Berman, B. P. et al. Regions of focal DNA hypermethylation and long-range hypomethylation in colorectal cancer coincide with nuclear lamina-associated domains. Nat. Genet. 44, 40–46 (2011).

51. Chen, S., Zhou, Y., Chen, Y. & Gu, J. fastp: an ultra-fast all-in-one FASTQ preprocessor. Bioinforma. Oxf. Engl. 34, i884–i890 (2018).

52. Li, H. et al. The Sequence Alignment/Map format and SAMtools. Bioinforma. Oxf. Engl. 25, 2078–2079 (2009).

53. Anders, S., Pyl, P. T. & Huber, W. HTSeq--a Python framework to work with high-throughput sequencing data. Bioinforma. Oxf. Engl. 31, 166–169 (2015).

54. Patro, R., Duggal, G., Love, M. I., Irizarry, R. A. & Kingsford, C. Salmon provides fast and bias-aware quantification of transcript expression. Nat. Methods 14, 417–419 (2017).

55. Ewels, P. A. et al. The nf-core framework for community-curated bioinformatics pipelines. Nat. Biotechnol. 38, 276–278 (2020).

56. Freese, N. H., Norris, D. C. & Loraine, A. E. Integrated genome browser: visual analytics platform for genomics. Bioinforma. Oxf. Engl. 32, 2089–2095 (2016).

57. Machanick, P. & Bailey, T. L. MEME-ChIP: motif analysis of large DNA datasets. Bioinforma. Oxf. Engl. 27, 1696–1697 (2011).

58. Ramírez, F. et al. deepTools2: a next generation web server for deep-sequencing data analysis. Nucleic Acids Res. 44, W160–165 (2016).

59. Krueger, F. & Andrews, S. R. Bismark: a flexible aligner and methylation caller for Bisulfite-Seq applications. Bioinforma. Oxf. Engl. 27, 1571–1572 (2011).

60. Ou, S. & Jiang, N. LTR_retriever: A Highly Accurate and Sensitive Program for Identification of Long Terminal Repeat Retrotransposons1[OPEN]. Plant Physiol. 176, 1410–1422 (2018).

61. Catoni, M., Tsang, J. M., Greco, A. P. & Zabet, N. R. DMRcaller: a versatile R/Bioconductor package for detection and visualization of differentially methylated regions in CpG and non-CpG contexts. Nucleic Acids Res. 46, e114 (2018).

62. Shimada, A. & Saze, H. RNA quality control by CCR4 safeguards chromatin integrity and centromere function in Arabidopsis. 2024.08.23.608929 Preprint at 10.1101/2024.08.23.608929 (2024).

