## Supplementary Materials (Tables and Figures) for "Nuclear envelope dysfunction drives premature aging and modulates heterochromatic methylome drift in *Arabidopsis*"

- Tables 1 to 3
- Supplementary Figures 1 to 14

| **Sample** | **Reads** | **Mapped reads** |
|---|---|---|
| col0-150DAS-1 | 74,832,580 | 44,001,557 |
| col0-150DAS-2 | 33,422,087 | 17,760,497 |
| *crwn1-1* | 31,350,791 | 17,609,739 |
| *crwn1-2* | 40,491,779 | 24,955,083 |
| *crwn2-1* | 38,482,774 | 22,127,595 |
| *crwn2-2* | 30,962,431 | 18,162,562 |
| *crwn3-1* | 28,827,278 | 16,595,864 |
| *crwn3-2* | 31,045,471 | 16,466,518 |
| *crwn4-1* | 81,117,845 | 51,923,533 |
| *crwn4-2* | 40,464,611 | 20,887,832 |
| *kaku4-1* | 33,777,264 | 21,195,233 |
| *kaku4-2* | 31,903,572 | 15,533,849 |
| col0-12DAS-1 | 54,290,054 | 51,092,370 |
| col0-12DAS-2 | 63,648,640 | 60,186,154 |

**Table 1 | Summary of RNA-seq sequencing and alignment metrics.**
For each sample, the table reports the total number of reads and mapped reads to the *Arabidopsis thaliana* TAIR10 reference genome.

| Sample | Reads (post filtering) | Mapped reads | Alignment rate (%) | Coverage (x) |
|---|---|---|---|---|
| 10DAS_IP_1 | 33,567,012 | 32,126,310 | 95.7 | 71 |
| 10DAS_IP_2 | 49,885,199 | 47,051,181 | 94.3 | 105 |
| 10DAS_IP_3 | 58,414,785 | 47,828,022 | 81.9 | 106 |
| 10DAS_INPUT_1 | 24,833,901 | 24,264,096 | 97.7 | 54 |
| 10DAS_INPUT_2 | 30,754,058 | 30,031,697 | 97.7 | 67 |
| 10DAS_INPUT_3 | 20,464,994 | 18,581,313 | 90.8 | 41 |
| 120DAS_IP_1 | 32,090,823 | 26,288,763 | 81.9 | 58 |
| 120DAS_IP_2 | 33,879,193 | 30,758,682 | 90.8 | 68 |
| 120DAS_IP_3 | 34,246,235 | 32,071,042 | 93.6 | 71 |
| 120DAS_INPUT_1 | 23,937,149 | 22,473,228 | 93.9 | 50 |
| 120DAS_INPUT_2 | 26,695,740 | 25,064,392 | 93.9 | 56 |
| 120DAS_INPUT_3 | 32,240,364 | 31,026,210 | 96.2 | 69 |
| untagged IP | 29,430,732 | 27,402,310 | 93.1 | 61 |
| untagged INPUT | 23,790,062 | 21,899,690 | 92.1 | 49 |

**Table 2 | Summary of KAKU4-eYFP ChIP-seq sequencing and alignment metrics.** For each sample, the table reports the total number of reads after filtering (Reads post-filtering), total number of reads mapped to the *Arabidopsis thaliana* TAIR10 reference genome (Mapped reads), alignment rate (Alignment rate [%]), and calculated sequencing depth (Coverage [×]) based on a 135 Mb genome size and a read length of 150 bp. "Reads" refers to individual mates from paired-end sequencing (R1 and R2 counted separately), as reported by samtools flagstat following adapter trimming and quality filtering.

| Sample | Reads (post deduplication) | Alignment rate (%) | Mapped reads | Coverage (x) | Conversion rate (%) | CG (%) | CHG (%) | CHH (%) |
|---|---|---|---|---|---|---|---|---|
| *col0-150DAS-1* | 31,996,098 | 82.3 | 26,332,789 | 59 | 99.8 | 26.9 | 10.9 | 5.7 |
| *col0-150DAS-2* | 30,148,076 | 77.8 | 23,455,203 | 52 | 99.9 | 26.6 | 11.0 | 6.0 |
| *crwn1-1* | 30,961,378 | 85.6 | 26,502,940 | 59 | 99.9 | 27.7 | 10.7 | 5.7 |
| *crwn1-2* | 29,576,109 | 82.9 | 24,518,594 | 54 | 99.9 | 27.9 | 11.1 | 5.7 |
| *crwn2-1* | 28,875,298 | 82.2 | 23,735,495 | 53 | 99.9 | 27.6 | 10.9 | 6.1 |
| *crwn2-2* | 58,439,744 | 83.1 | 48,563,427 | 108 | 99.9 | 27.8 | 11.3 | 6.2 |
| *crwn3-1* | 30,180,566 | 84.5 | 25,502,578 | 57 | 99.9 | 26.8 | 10.2 | 5.0 |
| *crwn3-2* | 27,319,983 | 85.0 | 23,221,986 | 52 | 99.9 | 26.7 | 10.3 | 5.1 |
| *crwn4-1* | 29,129,845 | 86.0 | 25,051,667 | 56 | 99.9 | 27.9 | 11.0 | 5.8 |
| *crwn4-2* | 22,842,258 | 82.1 | 18,753,494 | 42 | 99.9 | 27.5 | 10.8 | 5.6 |
| *kaku4-1* | 48,330,588 | 74.1 | 35,812,966 | 80 | 99.5 | 30.8 | 12.6 | 5.9 |
| *kaku4-2* | 48,566,688 | 83.0 | 40,310,351 | 90 | 99.9 | 24.9 | 9.1 | 3.7 |
| *crwn1crwn4-1* | 26,200,830 | 84.6 | 22,165,902 | 49 | 99.9 | 25.7 | 9.8 | 4.7 |
| *crwn1crwn4-2* | 46,763,274 | 83.5 | 39,047,334 | 87 | 99.8 | 25.9 | 10.0 | 4.7 |
| *col0-12DAS-1* | 28,977,191 | 84.3 | 24,427,772 | 54 | 99.9 | 26.4 | 9.9 | 3.5 |
| *col0-12DAS-2* | 23,159,077 | 79.9 | 18,504,103 | 41 | 99.8 | 26.8 | 10.8 | 4.1 |
| *kaku4-12DAS-1* | 27,605,631 | 81.4 | 22,470,984 | 50 | 99.9 | 26.9 | 10.4 | 4.2 |
| *kaku4-12DAS-2* | 24,875,555 | 77.9 | 19,378,057 | 43 | 99.8 | 27.8 | 10.8 | 4.0 |

**Table 3 | Summary of EM-seq sequencing and alignment metrics nuclear-envelope mutants.**

For each sample, the table reports the total number of reads after deduplication (Reads post-deduplication), alignment rate to the *Arabidopsis thaliana* TAIR10 reference genome (Alignment rate [%]), number of mapped reads (Mapped reads), and calculated sequencing depth (Coverage [×]) based on a 135 Mb genome size and a read length of 150 bp. Additional columns report bisulfite conversion efficiency (Conversion rate [%]) and the proportion of cytosines methylated in CG, CHG, and CHH contexts (CG [%], CHG [%], CHH [%]) across the genome.

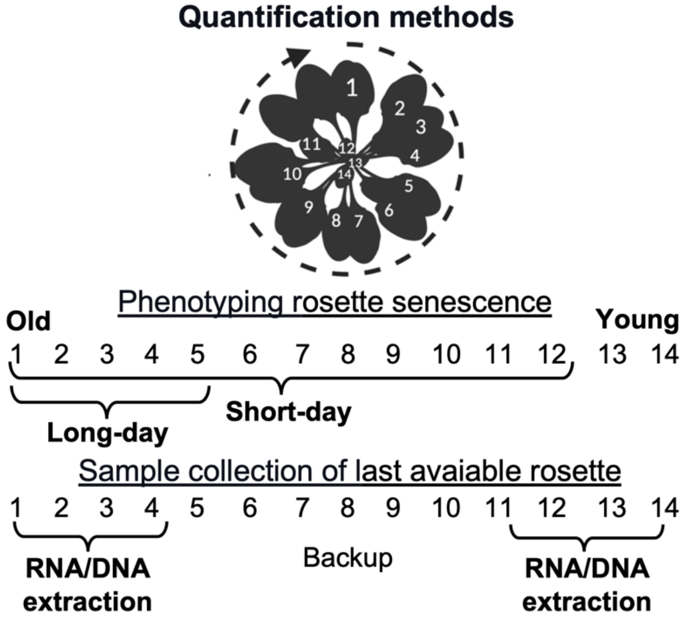


**Supplementary Figure 1 | Illustration: Standardized rosette phyllotaxy and sampling framework for spatial multiomics analyses.**

Leaves are numbered sequentially following the natural phyllotactic order (for simplicity represented in the circular arrow) from oldest (leaf 1) to youngest (leaf 14). Phenotyping of rosette senescence under LD conditions was based on the chlorosis and collapse of ≥5 oldest leaves; under SD conditions, ≥10-12 oldest leaves were assessed. For whole-rosette transcriptome and methylome profiling at 150 DAS (SD), basal leaf cohort (leaves 1-4) and distal leaf cohort (leaves 11-14) were pooled from three plants per replicate (n = 2 replicates per genotype), with a third pot retained as backup.

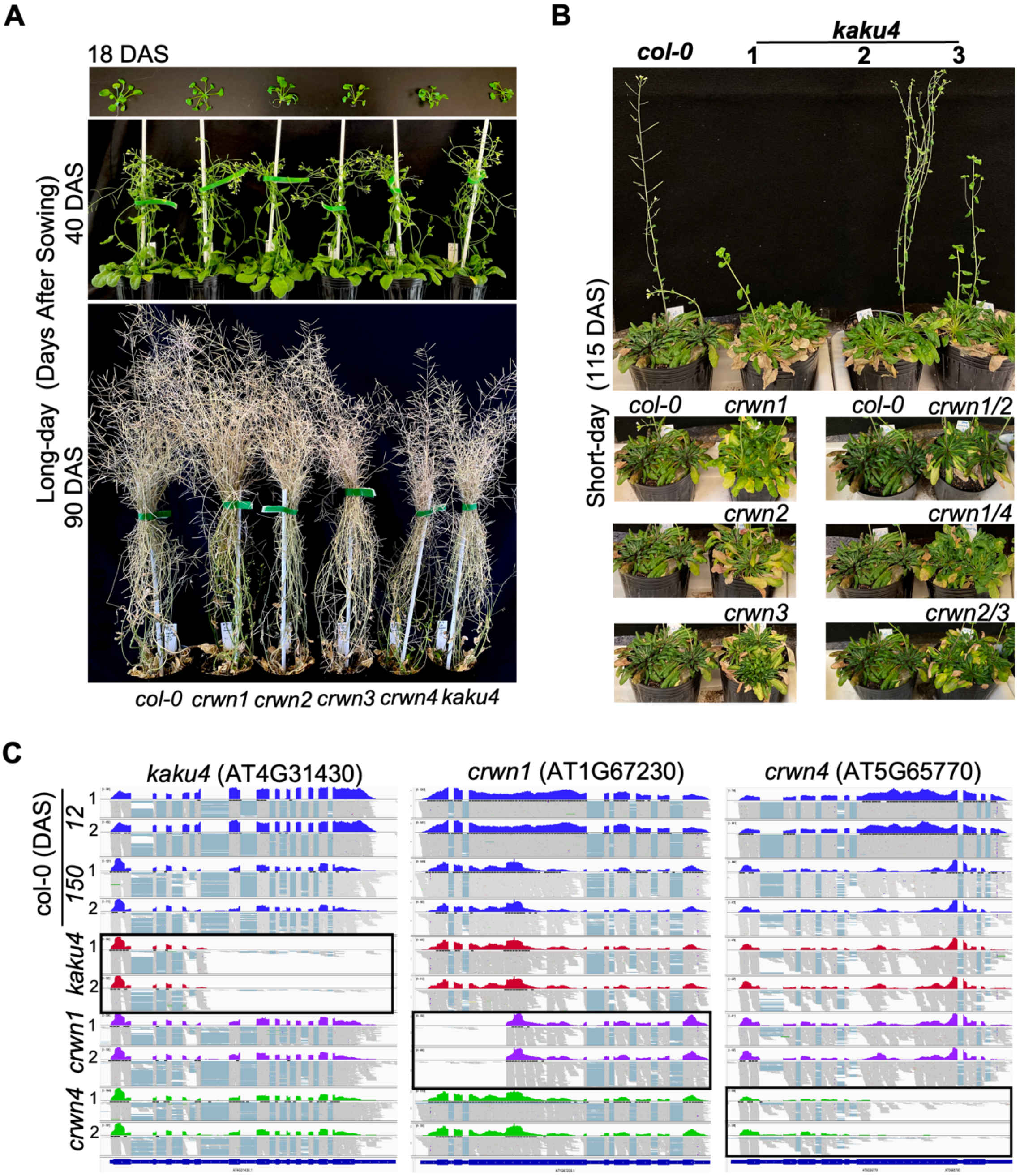


**Supplementary Figure 2 | Photoperiod modulation in LD does not affect lifespan of *Arabidopsis* nuclear-envelope mutants.**
**(A)** Time-course imaging (18, 40 and 90 DAS) of Col-0, *crwn1-4*, and *kaku4* mutants under LD. All genotypes successfully bolt and senesce by ~90 DAS.
**(B)** Phenotypes of *kaku4*, *crwn1-4* single and double mutants grown under SD conditions for 115 DAS. As in single mutations, double mutants also exhibit a progressive progeroid-like phenotype characterized by basal leaf necrosis, loss of meristematic competence, and bolting arrest.
**(C)** RNA-seq signal tracks at the *kaku4* (AT4G31430), *crwn1* (AT1G67230), and *crwn4* (AT5G65770) loci. Genome browser views of transcript abundance in wild-type (Col-0; 12 DAS and 150 DAS; blue), *kaku4* (red), *crwn1* (purple), and *crwn4* (green) mutants. Loss of full-length transcript accumulation at the corresponding locus in each T-DNA mutant confirms insertional disruption. Two biological replicates per genotype are shown.

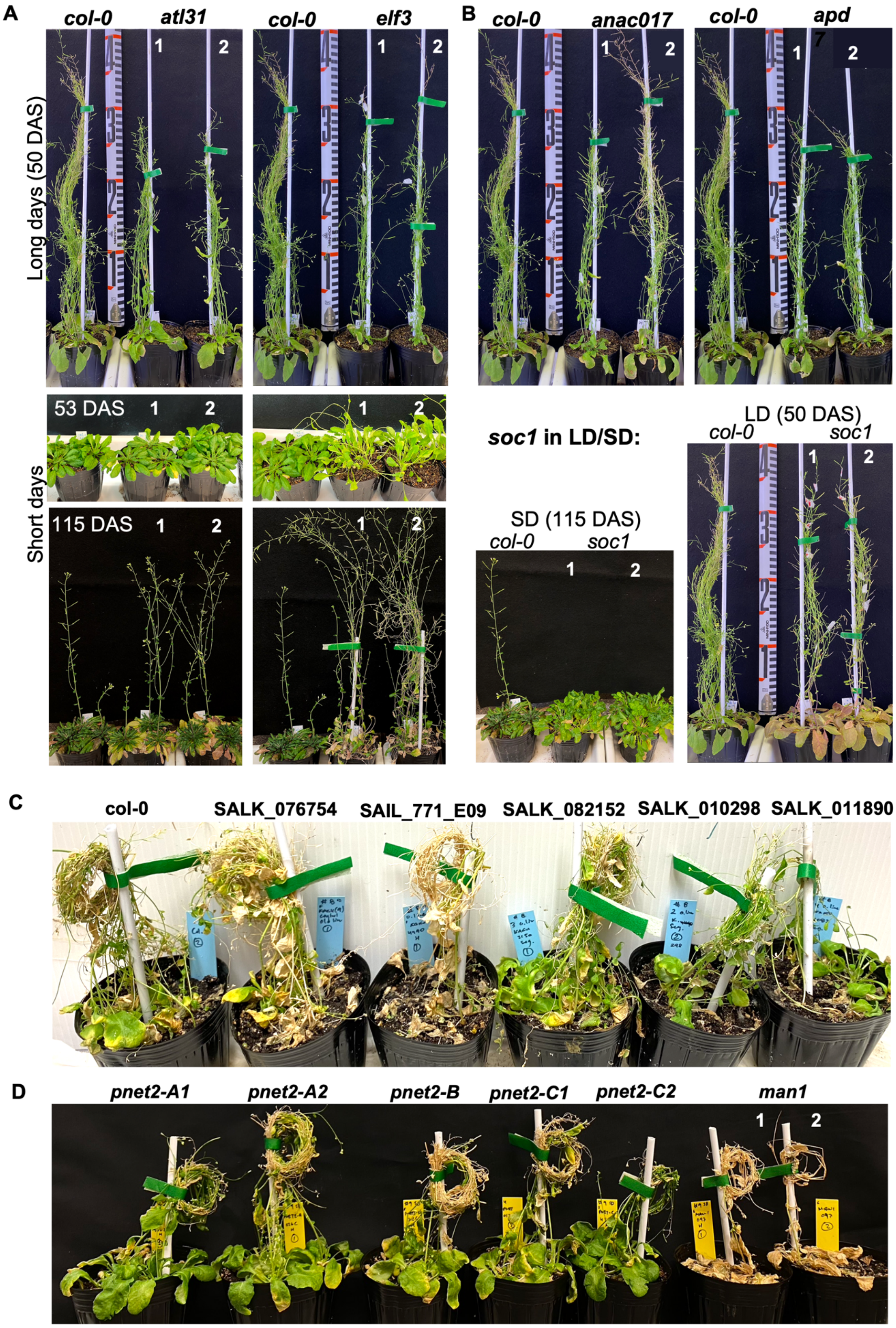


**Supplementary Figure 3 | Functional validation of photoperiod-response mutants for lifespan, flowering, and meristem fate in *Arabidopsis*.**

**(A)** Chronological lifespan and shoot elongation phenotypes of *ATL31* and *ELF3* mutants under long-day (LD) and short-day (SD) conditions. *ATL31* mutants, defective in a carbon/nitrogen-responsive E3 ubiquitin ligase, show accelerated senescence under SD at 115 DAS, consistent with loss of nutrient stress buffering. *ELF3*, a circadian evening complex component that represses *GI* and *CO*, exhibits photoperiod-independent early bolting and exaggerated shoot elongation under SD due to de-repression of floral integrators.
**(B)** Phenotypic characterization of *anac017*, *sppp/apd7*, and *soc1* mutants. Under LD (50 DAS), *anac017*—a NAC transcription factor regulating mitochondrial retrograde signaling—shows early reproductive transition and rosette decline. *sppp/apd7* double mutants (deficient in clade D PP2Cs that repress SAUR-mediated auxin output) exhibit severe vegetative collapse, implicating auxin misregulation in lifespan shortening. Under SD (115 DAS), *soc1* mutants fail to initiate bolting and maintain vegetative rosette identity, consistent with repression of the CO/FT/SOC1 module and a perennial-like apical arrest.
**(C)** Validation of *kaku4* loss-of-function across independent T-DNA alleles. Homozygous lines SALK_012298, SALK_082152, and SAIL_771_E09 (all targeting AT4G31430) were grown under SD for 130 DAS.
**(D)** Comparative analysis of *pnet2* and *man1* T-DNA insertion mutants under SD (120 DAS). *PNET2* single mutants retain vegetative integrity and bolting, while *man1* exhibit pronounced premature aging, loss of apical dominance, and inflorescence collapse.

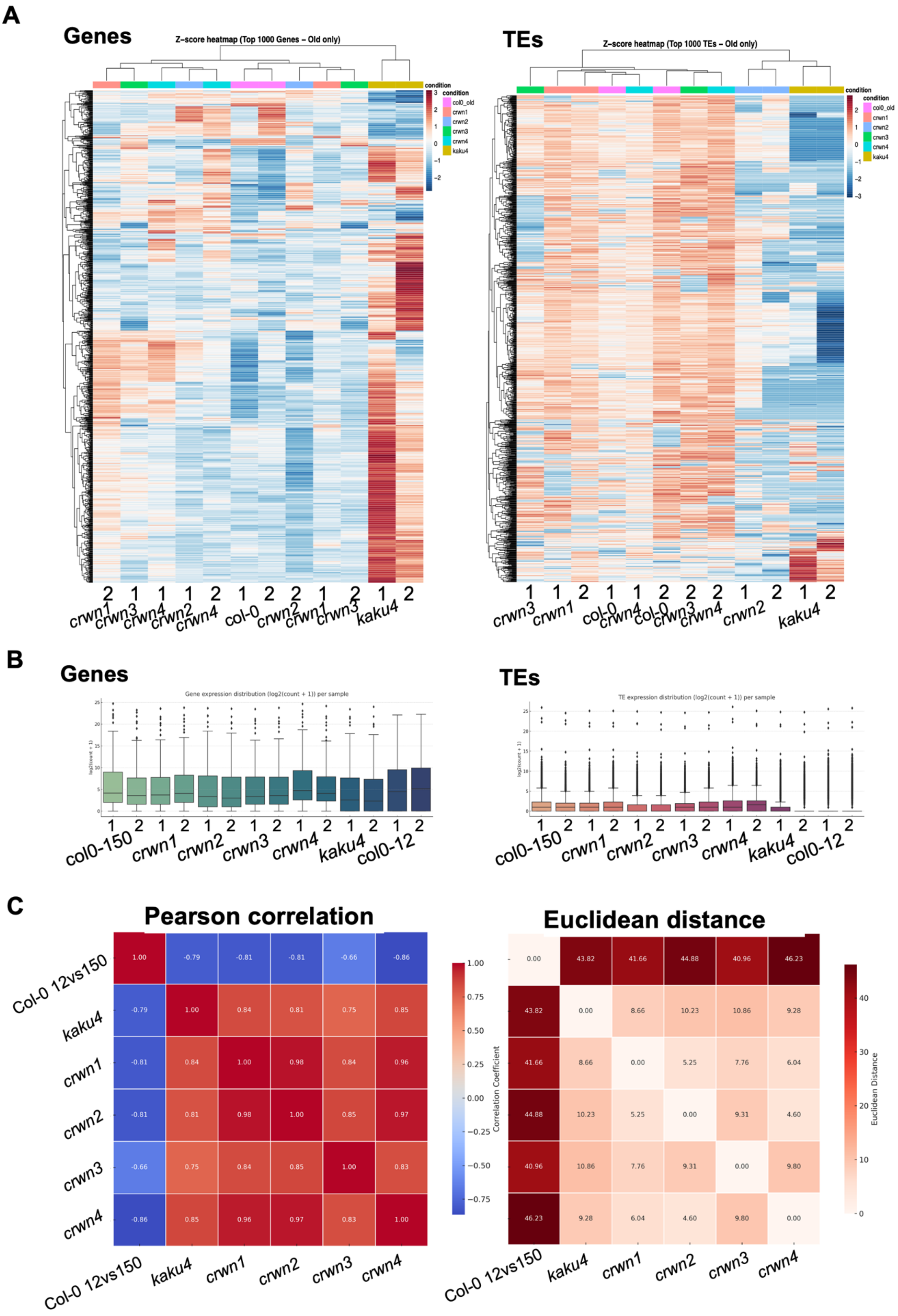


**Supplementary Figure 4. Expression distribution and variance-ranked Z-score heatmap of genes and TEs, and correlation matrix of expression profiles between libraries.**

**(A)** Variance-ranked Z-score heatmaps of gene and TE expression per genotype. Ranked by row-wise variance for 12 genotypes (150 DAS), and the top 1000 most variable features per category were selected. Rows were scaled by Z-score (mean-centered and unit-variance) and visualized using hierarchical clustering of both rows and columns.
**(B)** Distribution of protein-coding gene and TE expression across RNA-seq libraries. $Log_2$(count + 1) transformed expression values across 14 samples: 2 biological replicates each from wild-type (12 DAS and 150 DAS), *crwn1-4*, and *kaku4* mutants.
**(C)** Correlation and distance matrices among DEG profiles across aging and mutants. Left: Pearson correlation coefficients computed from $log_2$ fold-changes of DEGs shared between comparisons. Right: Euclidean distances between the same DEG $log_2$FC vectors. Diagonals represent self-comparisons.

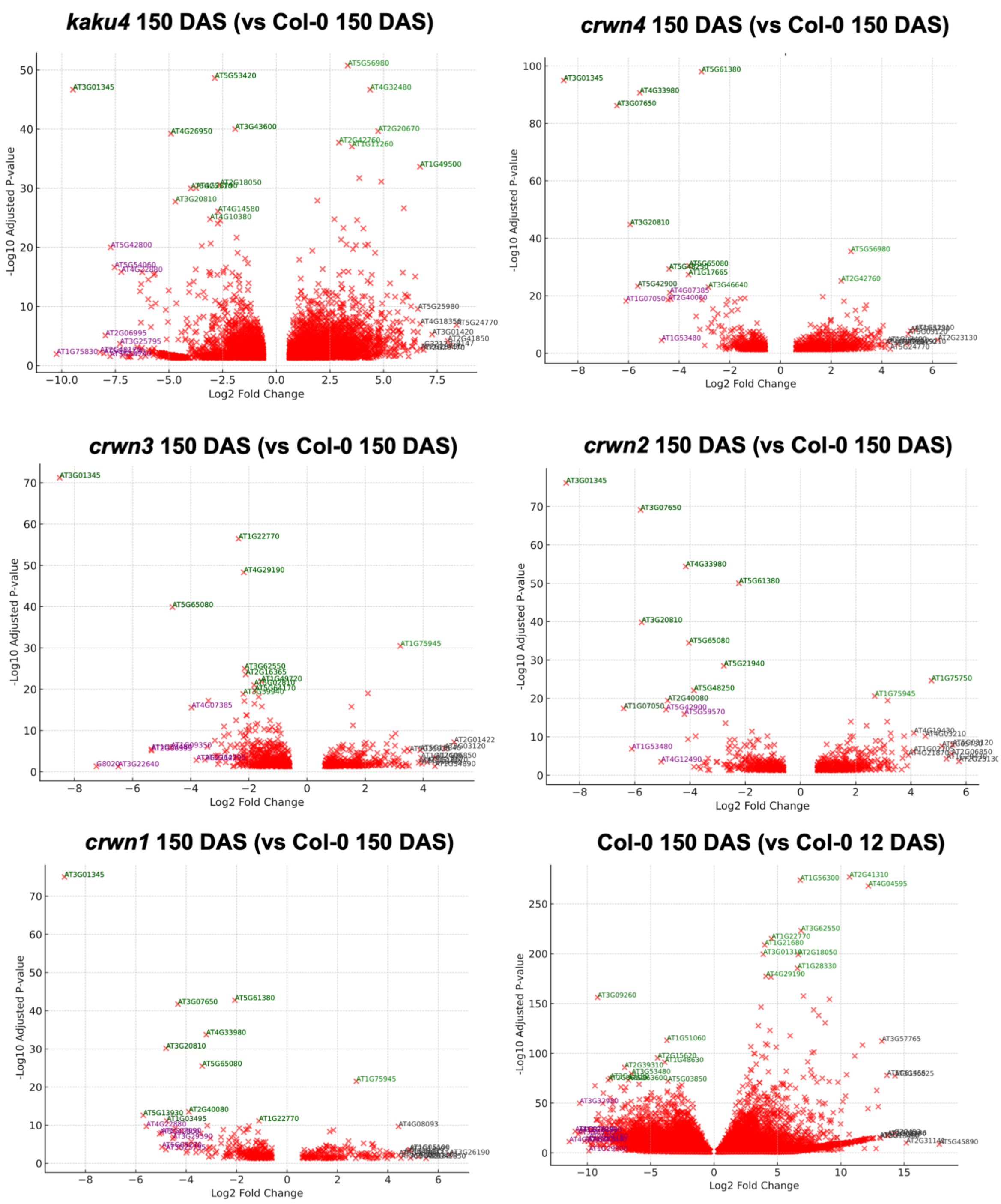


**Supplementary Figure 5 | Volcano plots for annotation enrichment during developmental aging and nuclear-envelope mutants.**
Volcano plots of $\log_2$ fold-change (x-axis) is plotted against $-\log_{10}$(adjusted p-value) (y-axis). Developmental aging is (Col-0 12 DAS vs 150 DAS) Mutants (at 150 DAS vs Col-0 150 DAS). Vertical dashed lines mark $\log_2$FC thresholds; the horizontal threshold indicates padj = 0.05 (mutants only). Top DEGs per contrast are labeled. Gene categories are color-coded based on prior annotation.

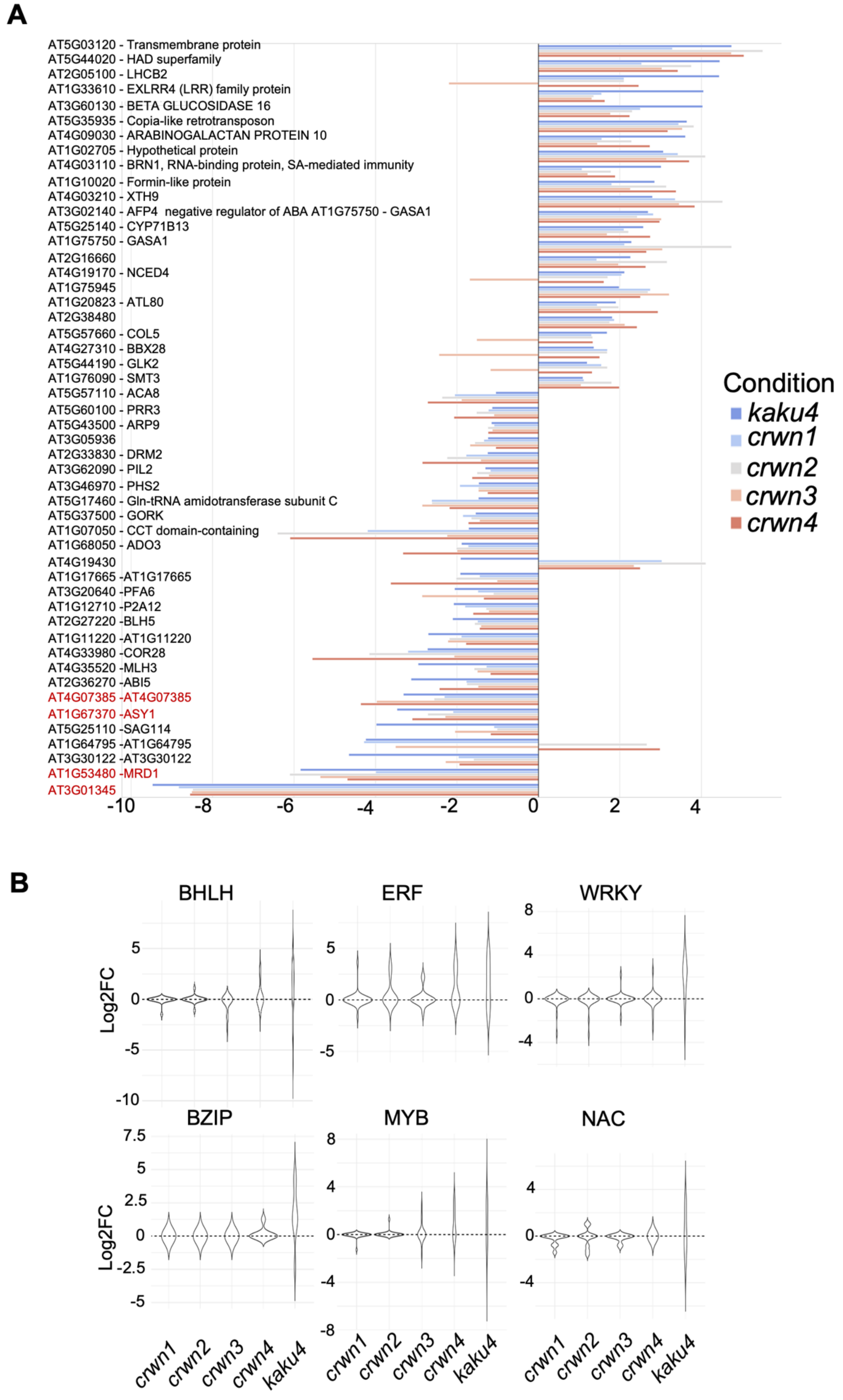


**Supplementary Figure 6 | Differentially expressed genes and gene families in nuclear-envelope mutants.**

**(A**) Horizontal bar plots show $log_2$ fold-change values of the top 25 up- and down-regulated transcripts shared between *kaku4* and *crwn1-4*. In red, lowest DEGs correspond to lncRNAs or 24-nt siRNAs islands, chromatin regulators, or senescence-linked loci (e.g., *MRD1*, *ASY1*).
**(B)** Differential gene expression across a *WRKY*, *NAC*, *ERF*, *MYB*, *BZIP*, and *SAG* genes, listed in **Fig. 3F,** showing the combined distribution of expression of genes per family in nuclear-envelope mutant (*kaku4*, *crwn1 to 4*).

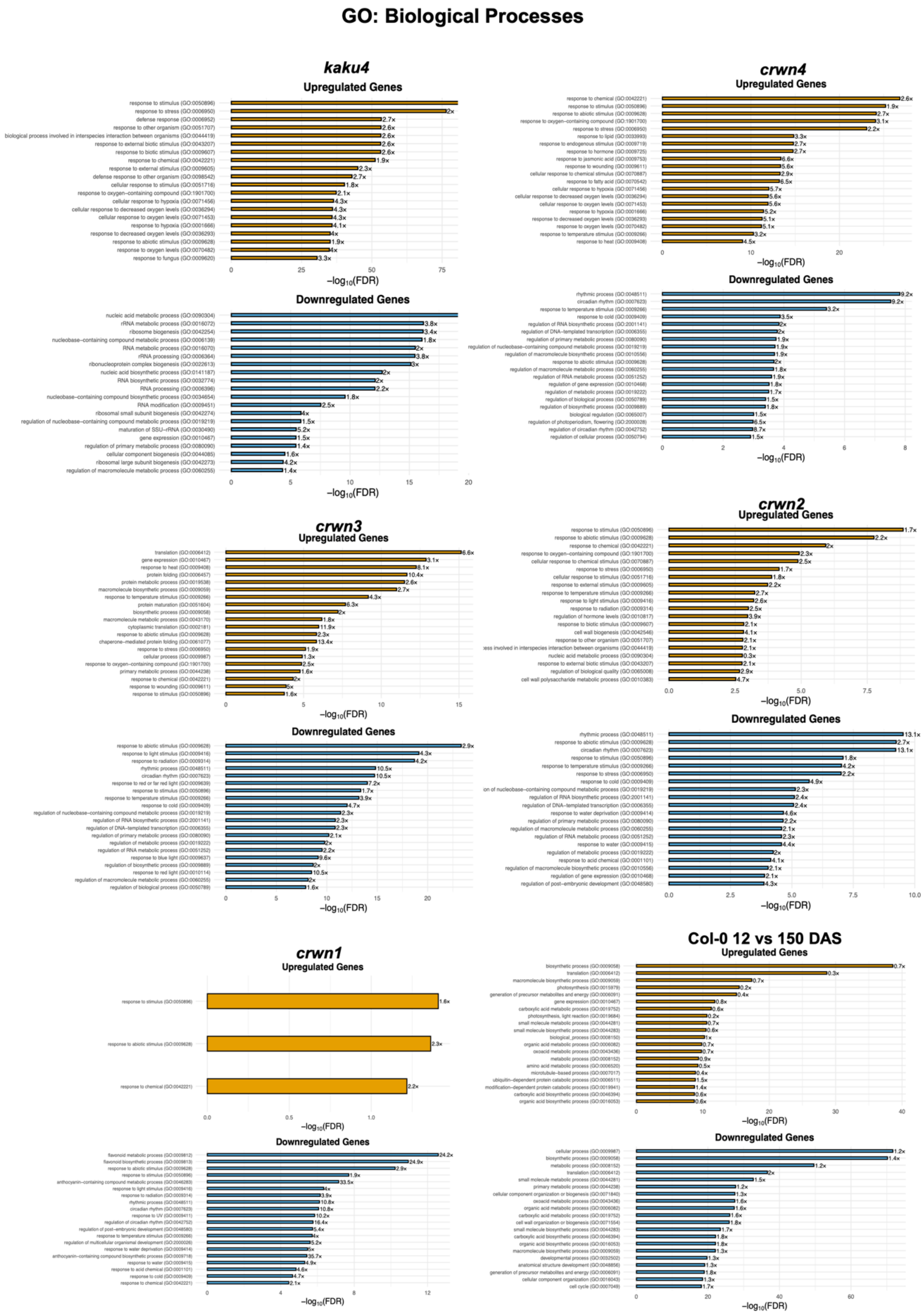


**Supplementary Figure 7 | Functional enrichment of DEGs for Biological Processes in developmental aging and during nuclear envelope dysfunction**. GO terms enriched

among upregulated (gold) and downregulated (blue) genes were identified using the PANTHER Overrepresentation Test with FDR correction and semantic similarity filtering (rrvgo). Bar plots show $-\log_{10}$(FDR) values and corresponding fold-enrichment for selected non-redundant categories from biological process, and molecular function, col0young correspond to Col-0 12 vs 150 DAS. See electronic version for higher resolution.

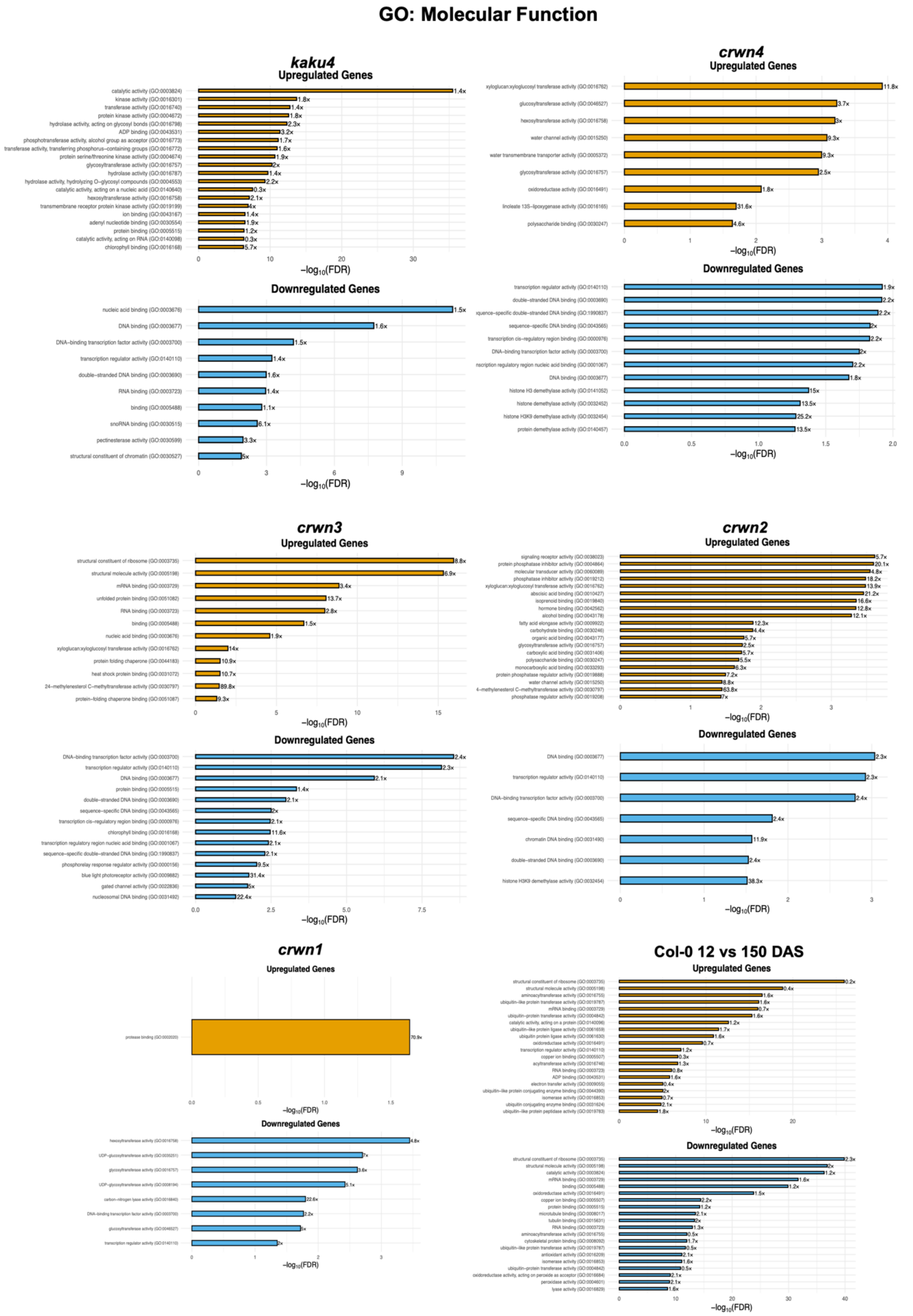


**Supplementary Figure 8 | Functional enrichment of DEGs for Molecular Functions in developmental aging and during nuclear envelope dysfunction.** GO terms enriched

among upregulated (gold) and downregulated (blue) genes were identified using the PANTHER Overrepresentation Test with FDR correction and semantic similarity filtering (rrvgo). Bar plots show $-\log_{10}$(FDR) values and corresponding fold-enrichment for selected non-redundant categories from biological process, and molecular function, col0young correspond to Col-0 12 vs 150 DAS. See electronic version for higher resolution.

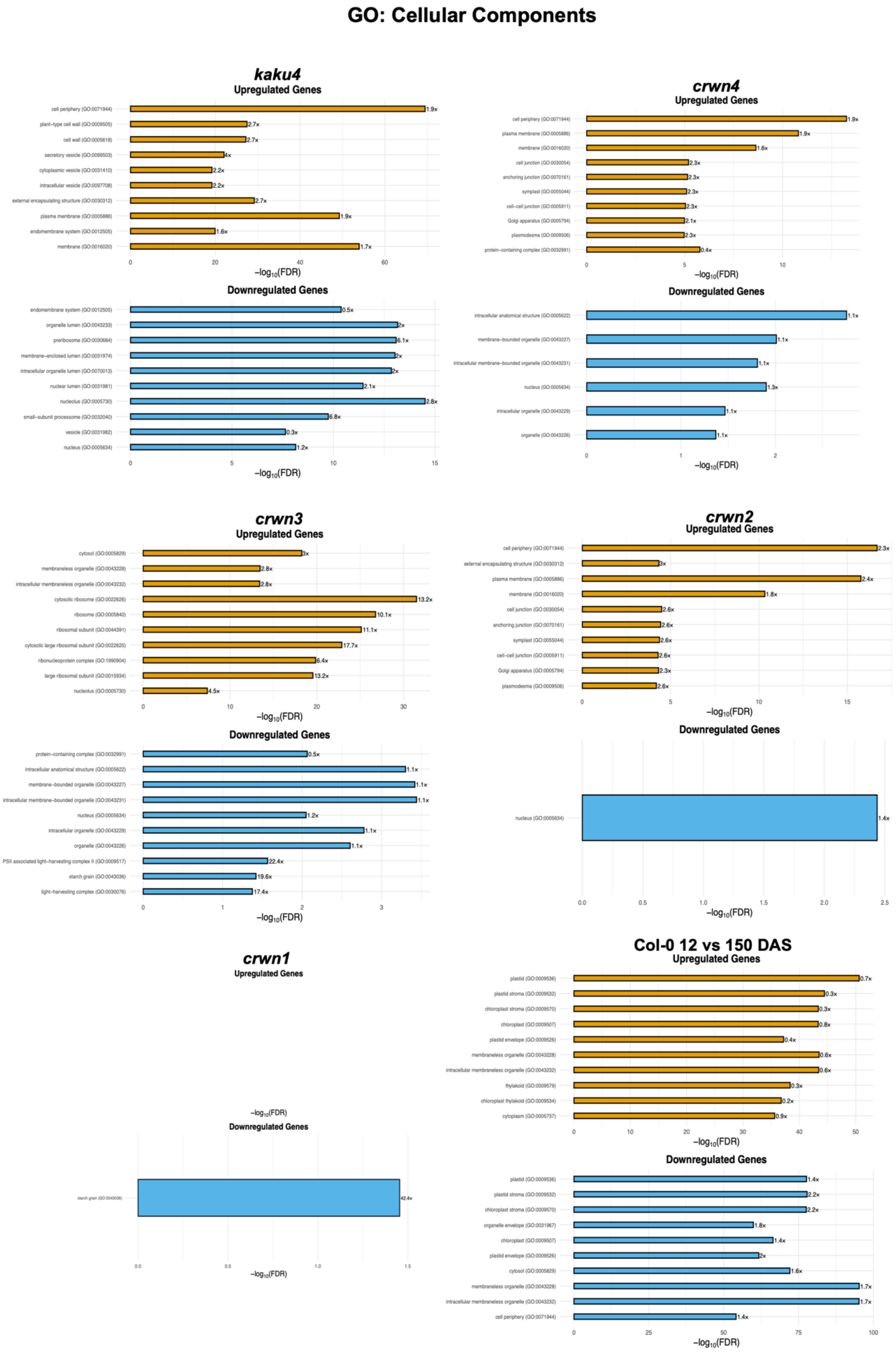


**Supplementary Figure 9 | Functional enrichment of DEGs for Cellular Components in developmental aging and during nuclear envelope dysfunction.** GO terms enriched

among upregulated (gold) and downregulated (blue) genes were identified using the PANTHER Overrepresentation Test with FDR correction and semantic similarity filtering (rrvgo). Bar plots show $-\log_{10}$(FDR) values and corresponding fold-enrichment for selected non-redundant categories from biological process, and molecular function, col0young correspond to Col-0 12 vs 150 DAS. See electronic version for higher resolution.

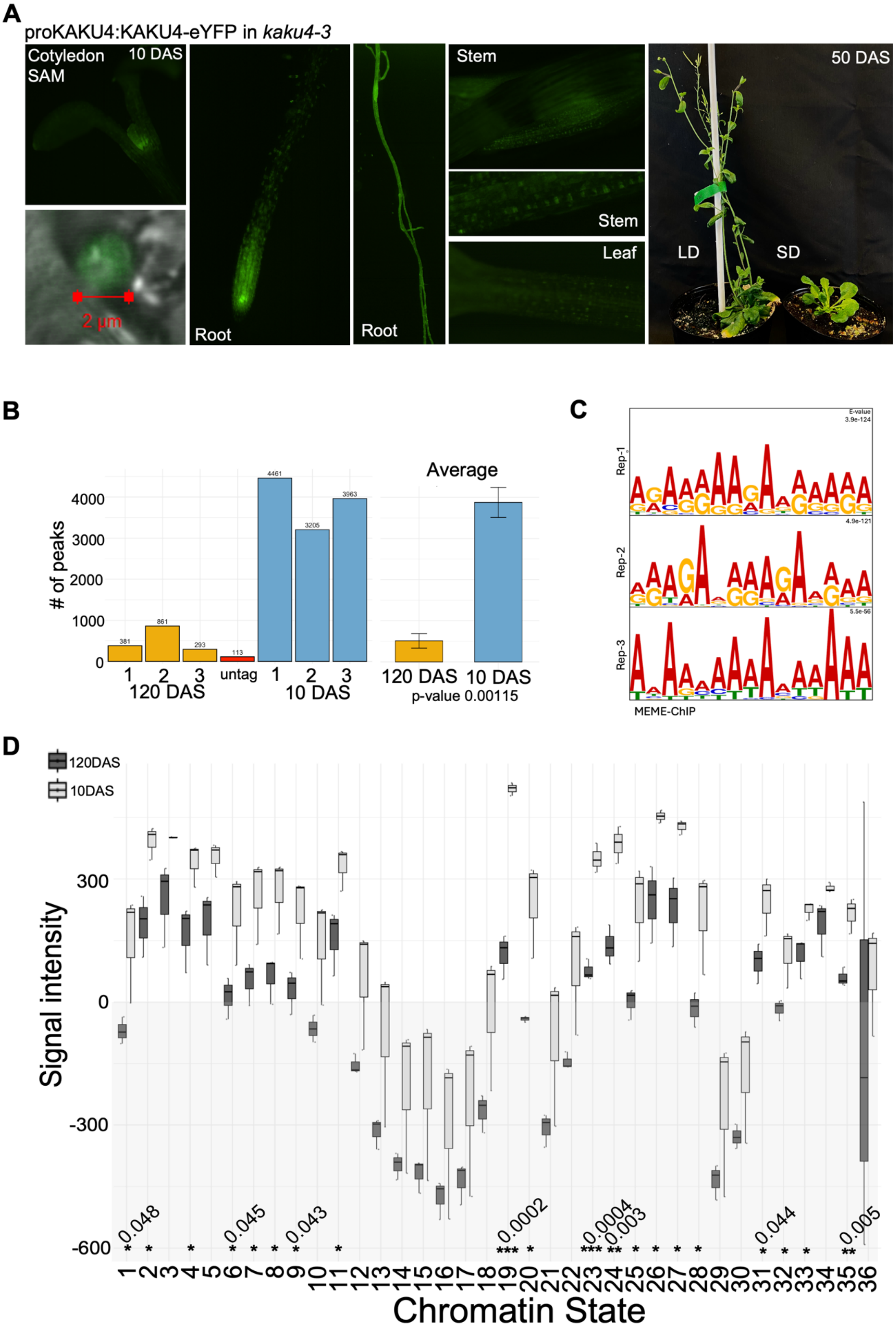


**Supplementary Figure 10 | KAKU4-eYFP reporter validation, global ChIP-seq peaks report and peak intensity across 36 Chromatin States of the *Arabidopsis* genome.**

**(A)** Subnuclear localization of KAKU4-eYFP fusion protein in 10 DAS seedlings. Expanded confocal images of proKAKU4:KAKU4-eYFP show perinuclear localization in cotyledon and root epidermal cells. Right: magnified nucleus highlights peripheral ring signal. Scale bar = 2 µm. Left: proKAKU4:KAKU4-eYFP in *kaku4-3* at 50 DAS in LD and SD showing normal development.
**(B)** Total number of MACS2-called broad peaks ($q < 0.05$, -broad-cutoff 0.1) in KAKU4-eYFP ChIP-seq from 10 DAS and 120 DAS tissues. Two-sided unpaired Welch's t-tests; bar charts show mean ± SD ($n = 3$).
**(C)** Motif enrichment from KAKU4-eYFP ChIP-seq peaks in young seedlings. Motif logos show top-ranked DNA motifs identified by MEME-ChIP from three independent 10-day-old replicates. All motifs are significantly enriched ($E < 1e-50$), with A/T-rich consensus signatures consistent across replicates. Input sequences were restricted to non-redundant peaks identified by MACS2 (broad mode, $q < 0.05$), normalized by input.
**(D)** Peak signal intensity at 36 chromatin states, calculated as the mean per-bin normalized coverage difference (IP - input) from ChIP-seq bigWigs, averaged across all peaks in that state for each replicate. Values are in arbitrary units from deepTools computeMatrix. Boxplots show $n = 3$; light grey = 10 DAS, black = 120 DAS. Two-sided t-test ($p < 0.05$) for group comparisons.

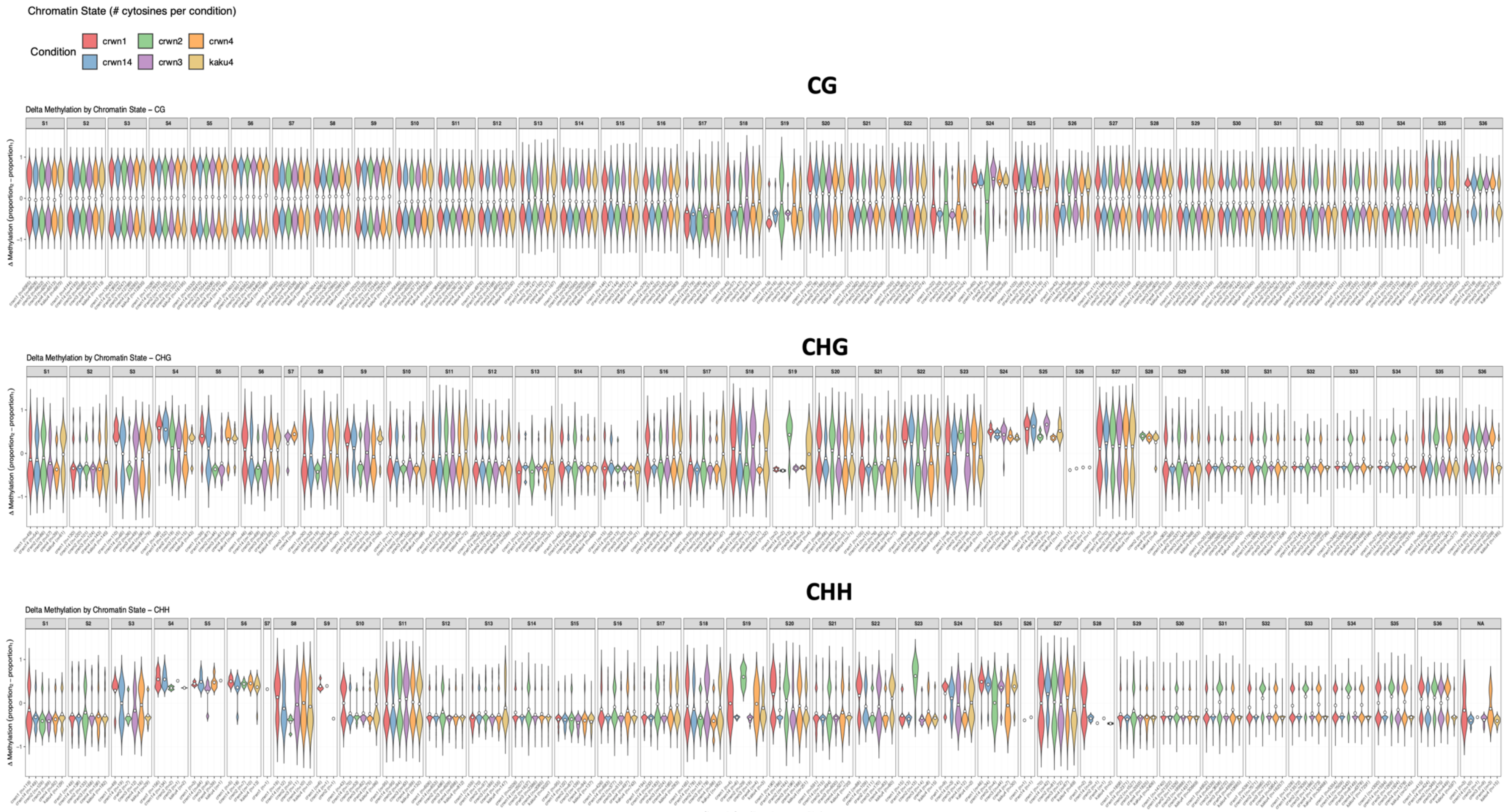


**Supplementary Figure 11 | Methylation drift in nuclear-envelope mutants across Chromatin States.**
Violin plots show the per-cytosine methylation difference (Δ methylation = mutant - WT 150 DAS) for all DMCs stratified by context (CG, CHG, CHH) across 36 Chromatin States. Each violin = a mutant, number of cytosines shown in x-axis. See electronic version for higher resolution. Violin plots for selected chromatin states in red boxes for Figure 4D.

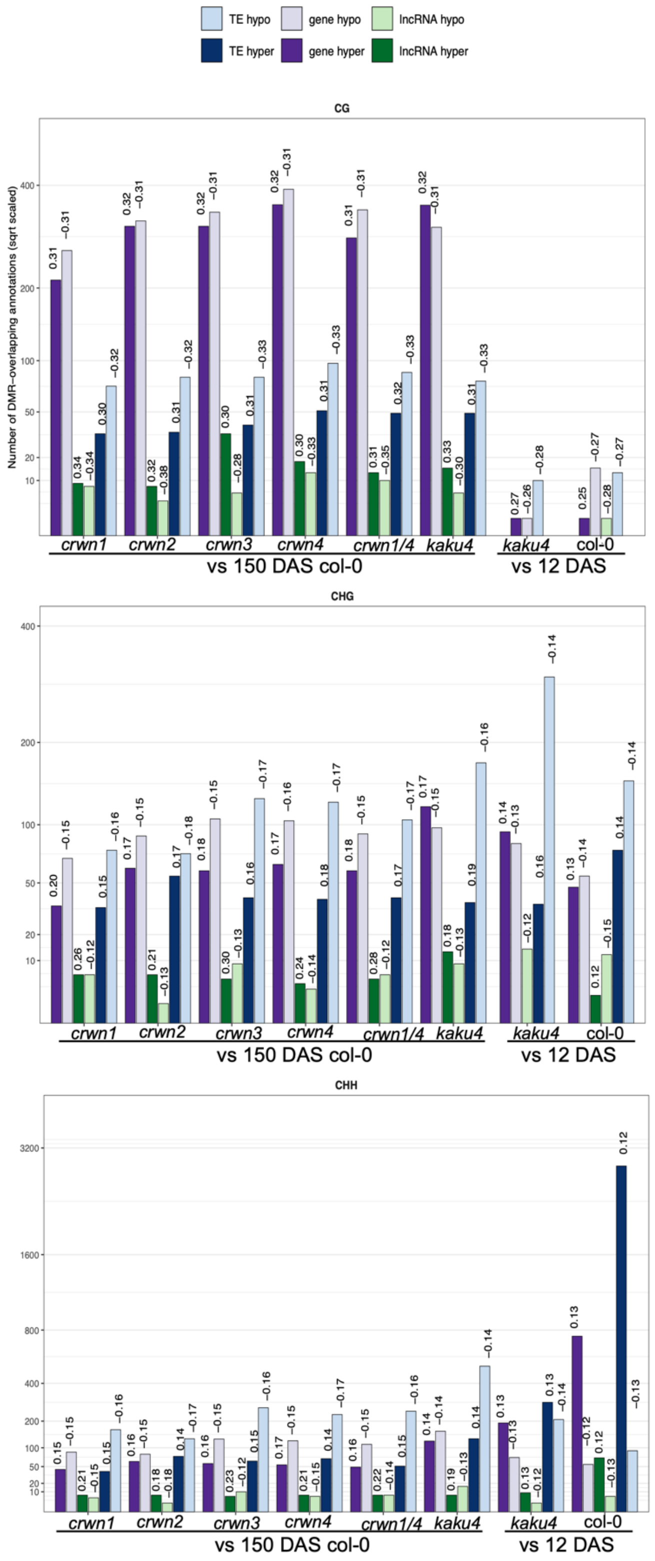


**Supplementary Figure 12 | Age- and mutant-associated DMRs at genes, TEs, lncRNAs; and multiomics annotations sorted by DMCs.**
Supporting Fig. 3.7C, annotation-resolved counts of CG/CHG DMRs overlapping TEs, protein-coding genes, and lncRNAs. DMRs were identified with thresholds (CG $\Delta m \geq 0.25$; CHG $\Delta m \geq 0.10$; CHH $\Delta m \geq 0.10$; FDR $q < 0.05$). Light bars denote hypomethylated and dark bars hypermethylated DMRs. Numbers above bars indicate the mean $\Delta m$ for each group.

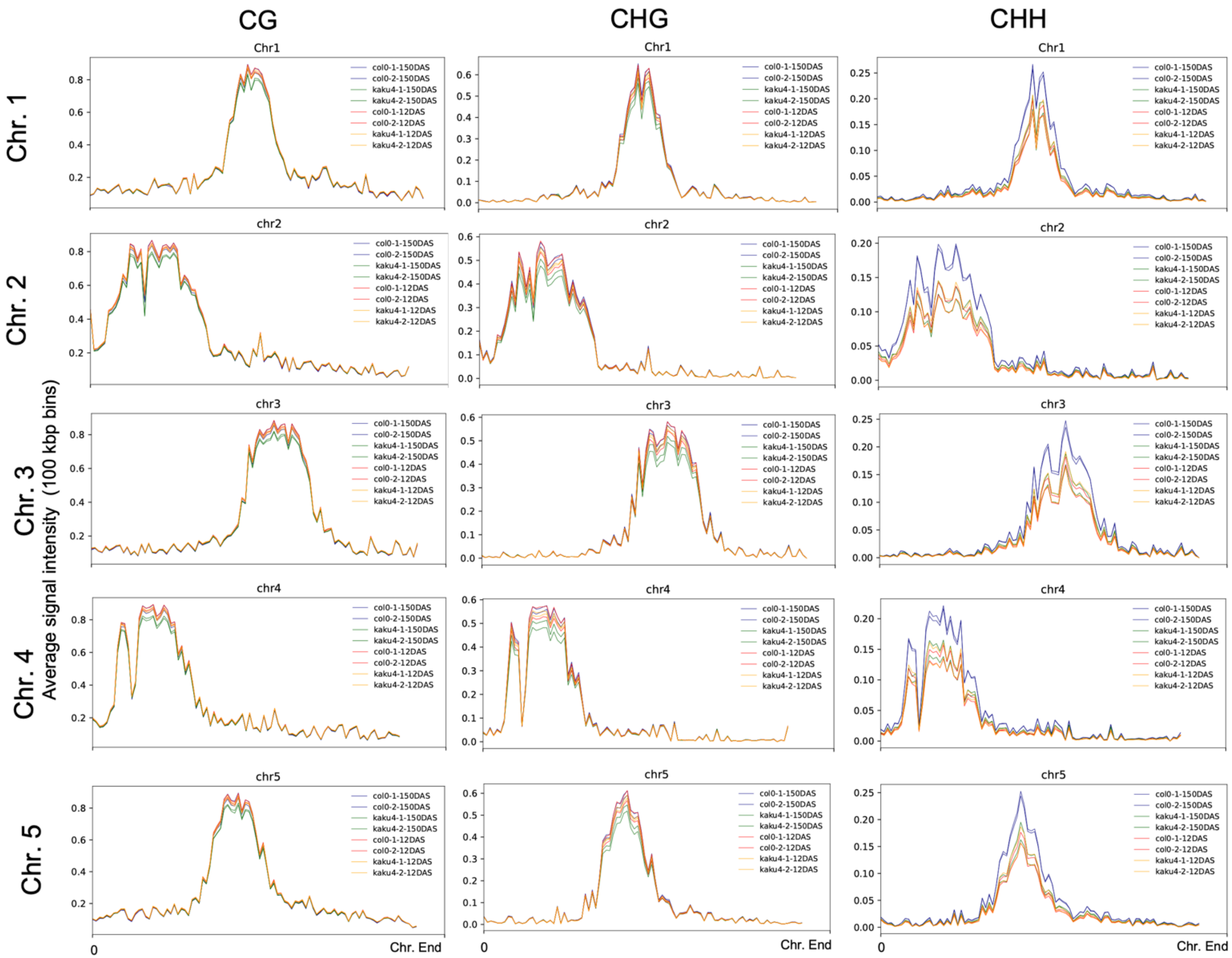


**Supplementary Figure 13 | Chromosome-scaled CG/CHG/CHH methylation profiles in wildtype and *kaku4* at 12 and 150 DAS.** Methylation profile along five chromosomes (binned in 100 kb windows). Col-0 and *kaku4* at 12 and 150 DAS, shown two replicates per condition.

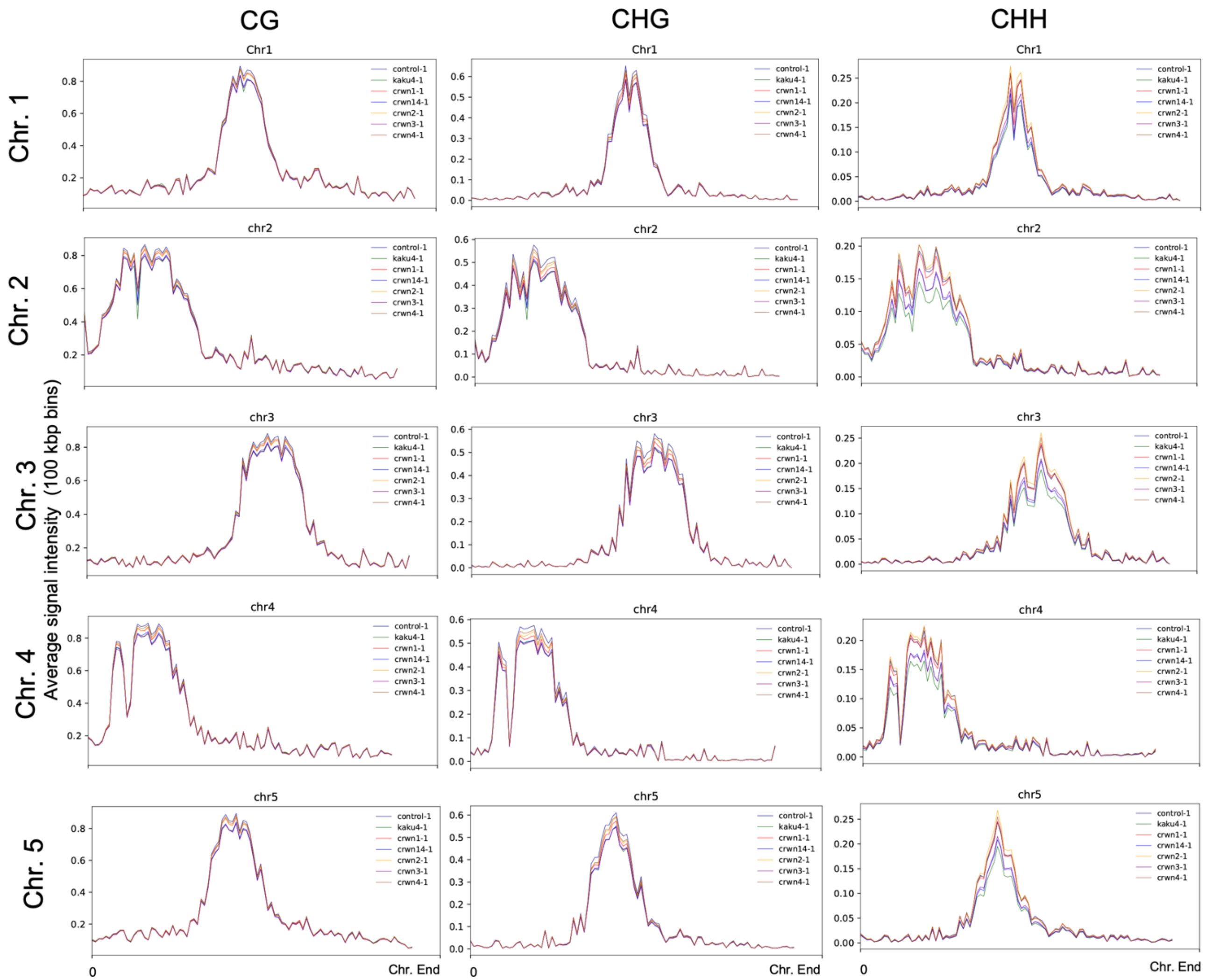


**Supplementary Figure 14 | Chromosome-scaled CG/CHG/CHH methylation profiles in wildtype and nuclear-envelope mutants.** Methylation profile along five chromosomes (binned in 100 kb windows). *crwn1-4* and *kaku4* at 150 DAS, shown one replicate for simplicity.
